# The *Arabidopsis thaliana* pan-NLRome

**DOI:** 10.1101/537001

**Authors:** Anna-Lena Van de Weyer, Freddy Monteiro, Oliver J. Furzer, Marc T. Nishimura, Volkan Cevik, Kamil Witek, Jonathan D.G. Jones, Jeffery L. Dangl, Detlef Weigel, Felix Bemm

## Abstract

Disease is both among the most important selection pressures in nature and among the main causes of yield loss in agriculture. In plants, resistance to disease is often conferred by Nucleotide-binding Leucine-rich Repeat (NLR) proteins. These proteins function as intracellular immune receptors that recognize pathogen proteins and their effects on the plant. Consistent with evolutionarily dynamic interactions between plants and pathogens, NLRs are known to be encoded by one of the most variable gene families in plants, but the true extent of intraspecific NLR diversity has been unclear. Here, we define the majority of the *Arabidopsis thaliana* species-wide “NLRome”. From NLR sequence enrichment and long-read sequencing of 65 diverse *A. thaliana* accessions, we infer that the pan-NLRome saturates with approximately 40 accessions. Despite the high diversity of NLRs, half of the pan-NLRome is present in most accessions. We chart the architectural diversity of NLR proteins, identify novel architectures, and quantify the selective forces that act on specific NLRs, domains, and positions. Our study provides a blueprint for defining the pan-NLRome of plant species.

## Introduction

Plant immune receptor repertoires have been shaped by millennia of plant-microbe coevolution^1,2^. Immunity is activated either by cell surface receptors that recognize microbe-associated molecular patterns (PAMPs), or by intracellular receptors that detect pathogen effectors^1^. These intracellular receptors are typically encoded by highly polymorphic genes. About two thirds of disease resistance genes encode nucleotide-binding leucine-rich repeat receptors (NLRs)^3^, and most plant genomes carry hundreds of NLR genes^4^. The majority of plant NLRs contain a central nucleotide binding domain shared between Apaf-1, Resistance proteins and CED4 (NB-ARC, hereafter NB for simplicity)^5^. Most contain also leucine-rich repeats (LRRs)^6,7^, and either a Toll/Interleukin-1 receptor (TIR) or coiled-coil (CC) domain at the N-terminus^8–10^. Proteins with similar arrangements of functional domains are also involved in host defense in animals and fungi^11–13^.

Recognition by NLRs generally involves one of three main mechanisms^14^. NLRs can directly detect pathogen effectors through interaction with the canonical NLR domains^15–17^, or with an NLR-incorporated integrated domain (ID) that resembles known domains of pathogen effector targets^18–22^. Alternatively, NLRs detect effector activity indirectly by monitoring a host virulence target (“guardee”)^23–25^, or detect effectors via direct interactions. Importantly, these mechanisms have been directly demonstrated only for a very small number of NLRs, and additional mechanisms might await discovery.

To date, NLR complements, or NLRomes, have been defined from available genome annotations for single cultivars of plants or for multiple species across different taxonomic levels, respectively^2,4,26–29^. The most striking findings were the repetitive modular arrangement of NLRs and the discovery of head-to-head paired NLR genes, of which one member included an ID^24,22,26–28^. The potential use of those IDs as modular building blocks has opened new possibilities for the engineering of novel resistances to pathogens^30–33^. The existing list of IDs, however, likely represents only a glimpse of the true diversity across plants.

The definition of pan-NLRomes, or repertoires of NLR genes, across different species, or higher taxonomic groups, has provided estimates of the variation in size of the NLR family^34–36^, presence/absence relations^35,36^, categorical distribution into structural classes across the phylogeny, and diversity of IDs^26,27^. Publicly available plant genome annotations have been the foundation of most NLRome studies and, their systematic integration has allowed ancestry reconstruction of key NLR lineages and illuminated ancient and recent expansion-contraction events^4^. In contrast, knowledge of the true diversity of within species pan-NLRomes is scarce and has so far been derived from only a limited number of individuals, and thus covers a narrow diversity within the population^34–37^. Across individuals of the same species, which often has only a single reference genome annotation, the remarkable differences in NLR family size between rice, tomato, and *Arabidopsis thaliana* might be due to low coverage of available genomes, or the difficulty of accurate assembly of tandem paralogous genes often found in NLR clusters when short-read sequencing is used under conditions of insufficient depth^34,35^.

Despite these potential shortcomings, early intraspecific pan-NLRome studies revealed patterns of allelic and structural variation consistent with adaptive evolution and balancing selection for subsets of NLR-encoding genes^37^, fitting a model of co-evolution of host and pathogens. Allelic variation seems to be reflected in many different haplotypes that are found across NLR loci^38,39^. These can include recombination “hotspots” generating NLR clusters^36,40^, and true allelic series^15,41^. The patterns of presence/absence polymorphisms as well as copy number variation at loci with multiple NLR genes imply that reference genomes may not include representatives of all distinct NLR clades within a species^36,38,42–44^. *<u>R</u>esistance* gene <u>en</u>richment <u>seq</u>uencing (RenSeq) facilitates discovery of “missing” NLR genes in a species, especially when hybridization-based capture of genomic fragments with sequence similarity to known NLR-coding genes is combined with <u>S</u>ingle-<u>M</u>olecule <u>R</u>eal <u>T</u>ime sequencing (SMRT RenSeq)^45^.

Our objective was to define the full NLR repertoire and its variability in the reference species *A. thaliana*, by analyzing a panel of 65 diverse accessions using SMRT RenSeq. We show that we reach saturation of the pan-NLRome with this well-chosen set of accessions; we define the core NLR complement of the species and detail novel domain architectures; and we describe presence-absence polymorphisms in non-core NLRs. Together, our work provides a foundation for the identification and cloning of disease resistance genes in more complex species of agronomic importance.

## Results

### The Samples

A set of 65 *A. thaliana* accessions was selected to explore the diversity of the pan-NLRome (Fig. 1a, Supplementary Table 1). The selection included 46 accessions from the 1001 Genomes Project, of which 21 belonged to previously identified relict populations characterized by an unusually high amount of genetic diversity^46^. Additionally, the 19 founder accessions of MAGIC lines, a resource to dissect the genetics of complex traits^47,48^ and the widely-studied accession Ws-2, were included.

**Figure 1.**
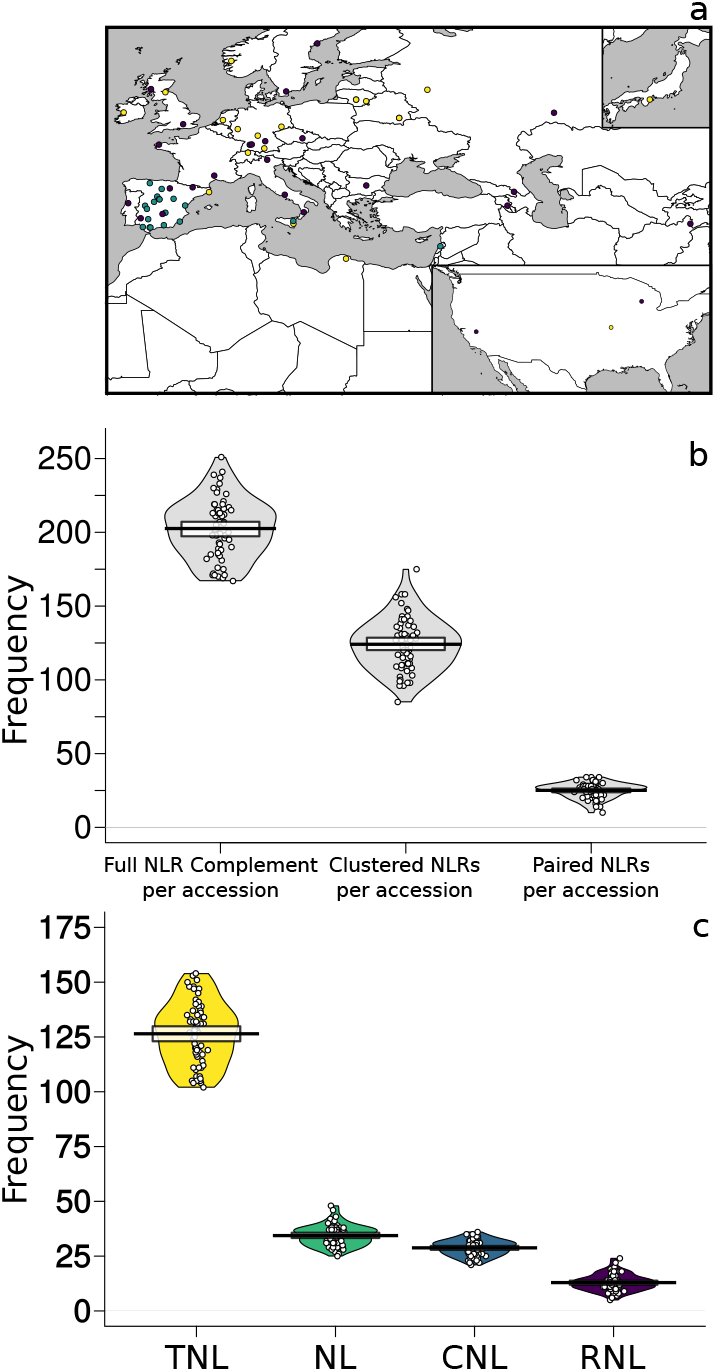
Overview of NLR complements in 65 accessions. a) Map of accession provenance. 1001 Genomes (relicts, blue, non-relicts, purple), MAGIC founders (yellow). b) Number of total, clustered and paired NLRs in accessions. Means, solid black lines; Bayesian 95% Highest Density Intervals (HDIs), solid bands. Individual data, open circles; full densities shown as bean plots. c) Number of different structural classes in accessions. Mean, HDI, and individual data as in (b).

### NLR Complements

Baits were designed to hybridize with 736 NLR-coding genes from multiple Brassicaceae, including *Arabidopsis thaliana, Arabidopsis lyrata, Brassica rapa, Aethionema arabicum* and *Eutrema parvulum* (see Online Methods for details of bait design, sequencing, assembly, annotation and quality control approaches). A combination of NLR sequence capture (RenSeq) and single-molecule real-time sequencing (SMRT) was used to reconstruct full NLR complements. In total, we produced 13,167 NLR-coding gene annotations, with a range of 167 to 251 genes per accession (Fig. 1b). Individual accessions had between 47% and 71% physically clustered NLR genes (more than one NLR in 200 kb of genomic sequence; adapted from^49^. A particularly interesting subset of NLR-coding genes are those in head-to-head orientation^50,51^, and we found 10 to 34 NLRs per accession in such an orientation, or with high sequence similarity to known functional pairs (see Online Methods). NLRs were grouped into four classes (TNL, NL, CNL, and RNL; see Glossary) based on canonical protein domains (TIR, NB, CC, RPW8 and LRR). Across all accessions, TNLs formed the largest and most size-variable class, followed by NLs, CNLs, and RNLs (Fig. 1c, Supplementary Fig. 1). Of the 13,167 NLR genes, 663 contained at least one additional integrated protein domain (ID), in which we found 36 distinct Pfam domains (Fig. 2a,b, Supplementary Table 2 and Supplementary Table 3). Individual accessions had 5 to 17 IDs distributed across 4 to 16 NLR genes, in line with reports for specific accessions^4,36^. This result reveals an unprecedented incidence of previously unreported *A. thaliana* IDs. Annotated RenSeq Col-0 identifiers and sequences are provided as supplementary material, but the Col-0 TAIR10/Araport11 sequences and identifiers were used in all downstream analysis (see Online Methods).

**Figure 2.**
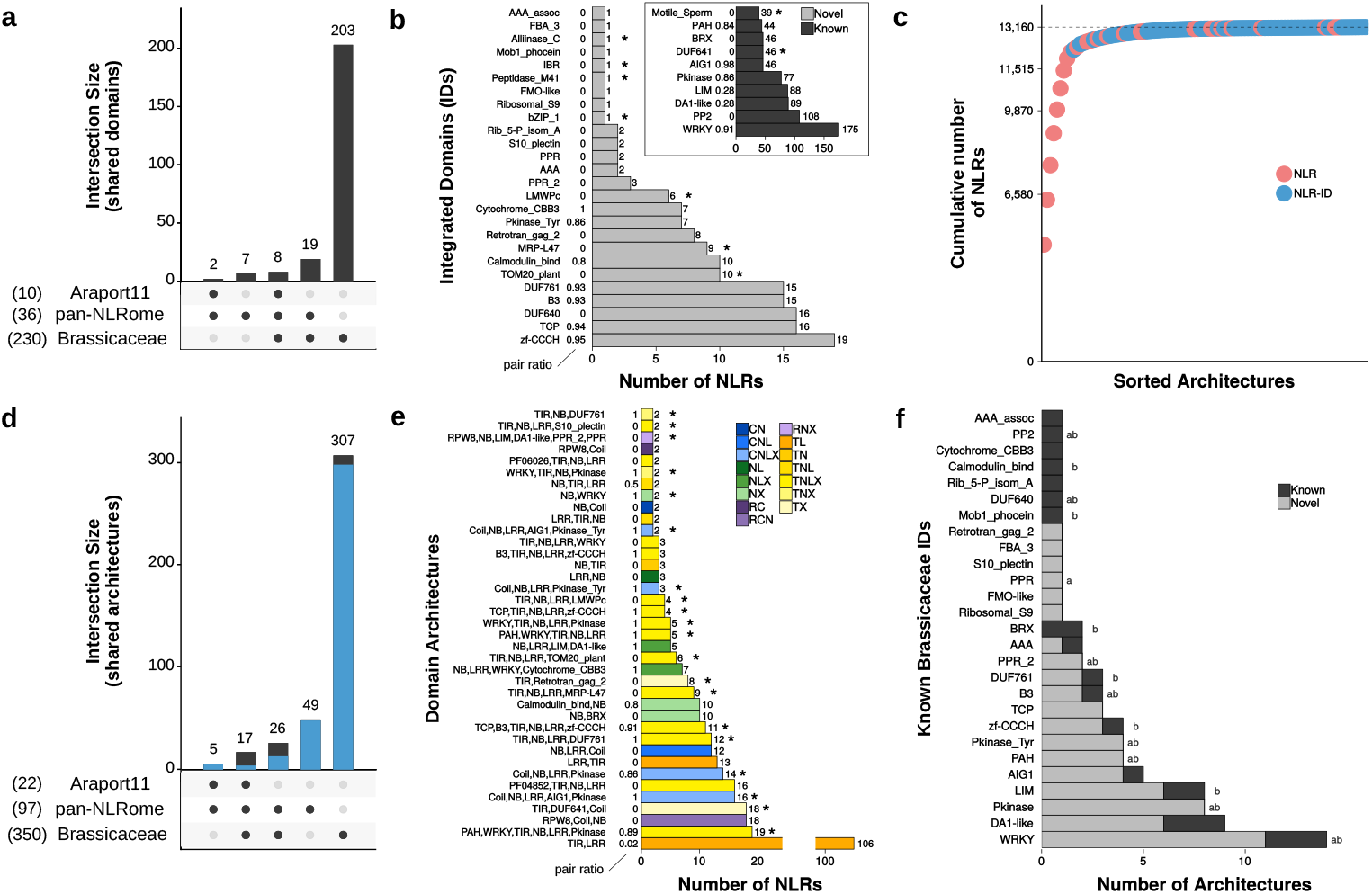
Diversity of IDs and domain architectures. a) UpSet intersection of IDs in the pan-NLRome, the Col-0 reference accession and 19 other Brassicaceae. The number of IDs in each set is indicated between parentheses at the lower left. Set intersections shown on the bottom. b) ID analysis. IDs known for *A. thaliana* in black and IDs newly described for *A. thaliana* in light grey. Asterisks indicate IDs not found in other Brassicaceae. Numbers next to y-axis indicate ratio of paired NLRs among ID containing NLRs. c) Cumulative size contribution to the pan-NLRome (y-axis) of each of the 97 size sorted domain architectures (x-axis). d) UpSet intersection plot of architectures shared between pan-NLRome, *A. thaliana* reference- and 19 other Brassicaceae. e) 38 architectures with at least two representatives and not found in the Col-0 reference. Asterisks indicate 20 architectures not found in 19 Brassicaceae family genomes or the *A. thaliana* Col-0 reference. Numbers next to y-axis shows fraction of paired NLRs. f) Number of known and novel architectures containing the 27 overlapping Brassicaceae IDs (see a). “*a*”and “*b*” to the right of the bars indicate putative IDs^26,27^ (see also Supplementary Table 2).

### NLR Domain Architecture Diversity

We investigated the repertoire of the 36 IDs, since these might function as pathogen effector binding platforms^19–21,32^. 29 of the 36 IDs were already known from other Brassicaceae including *A. thaliana* Col-0 (Fig. 2a, b; Supplementary Table 2, Supplementary Table 4, Supplementary Table 5, Supplementary Table 6). Nine of the 36 IDs were reported concordantly in the two major NLR-ID censuses, namely WRKY, PP2, Pkinase, PAH, DUF640, B3, Pkinase_Tyr, PPR_2 and Alliinase_C^26,27^. Five of those nine occur in genetically linked paired NLRs in the pan-NLRome (pair ratio > 0.5 in Fig. 2b, see Online Methods and Glossary; Supplementary Table 2). Rediscovery of these nine IDs is of relevance, since these are enriched for domains similar to known effector targets^26,27,52,53^. Our sequencing and annotation effort expands the *A. thaliana* ID repertoire beyond the ten IDs found in the Col-0 reference accession. IDs found in only one gene model did not receive particular attention, as they are conceivably an artefact of our annotation pipeline.

A hallmark of NLRome variation across species is the variation in the relative fraction of different domain architectures^4,11^. Examining the arrangement of NLR domains in the *A. thaliana* pan-NLRome we identified 97 distinct architectures (Supplementary Fig. 2). Whilst 27 canonical architectures (without IDs) account for the vast majority of the identified NLRs (95% of the pan-NLRome), the remaining 5% contain at least one of 36 different IDs (Fig. 2c). The 97 architectures greatly augment the 22 architectures found in the reference Col-0 genome (Fig. 2d, Supplementary Table 7), with most of the new *A. thaliana* architectures containing at least one ID (Supplementary Fig. 2). Half of the new *A. thaliana* architectures contain more than one gene (38/75) (Fig. 2e), of which, 17 predominantly comprise paired NLRs (pair ratio > 0.5, see Online Methods and Glossary) and contain at least one ID (Fig. 2e). About half of the architectures have not been previously described in the Brassicaceae (including *A. thaliana* Col-0) (49/97) (Fig. 2d, Supplementary Table 7). These novel architectures account only for 1.3% of the pan-NLRome (175 NLRs), with all but one containing an ID (Fig. 2d, e, Supplementary Table 7, Supplementary Table 8 and Supplementary Table 9). Finally, 12 IDs are repeatedly recruited into different novel architectures (labeled “novel > known” in Fig. 2f, Supplementary Table 8 and Supplementary Table 9), reflecting the recycling of a limited set of IDs into new domain arrangements. It is likely that these IDs are derived from proteins repeatedly targeted by pathogen virulence effectors.

### The pan-NLRome

To begin to understand the diversity of both NLR content and alleles, we grouped sets of homology-related NLRs from different ecotypes. The resulting clusters were termed orthogroups. We clustered 11,497 NLRs into 464 high confidence orthogroups (OGs) (Fig. 3a), plus 1,663 singletons. Ninety-five percent of the OGs could be discovered within 38 randomly chosen accessions (Fig. 3b). Additional sampling only recovered OGs with three or fewer members, indicating that the pan-NLRome we describe is largely, if not completely, saturated. OGs were classified according to size, domain architecture and structural features. We define the core NLRome as the 106 OGs found in at least 52 accessions (6,080 genes), 143 OGs found in at least 13, but fewer than 52 accessions as shell (3,932 genes), and the 215 OGs found in 12 or fewer accessions as cloud (1,485 genes) (Fig. 3a). The majority of OGs, 58%, were TNLs, in concordance with TNLs being the prevalent NLR class in the *Brassicaceae*^54^, 22% were CNLs, 7% RNLs, and 13% NLs (Fig. 3c). TNLs showed a strong tendency towards larger shell and core OGs compared to CNLs (Supplementary Fig. 3). Sixty-four OGs included genetically paired NLRs (see Online Methods), and 28 contained members with an ID, with almost none being present in the cloud NLRome (Fig. 3d). Shell and core OGs contained most paired NLRs (98% in 55 OGs, Supplementary Fig. 3). This shows that conserved NLR pairs are widely distributed in the population and that incorporation of IDs into NLRs is widespread in *A. thaliana*.

**Figure 3.**
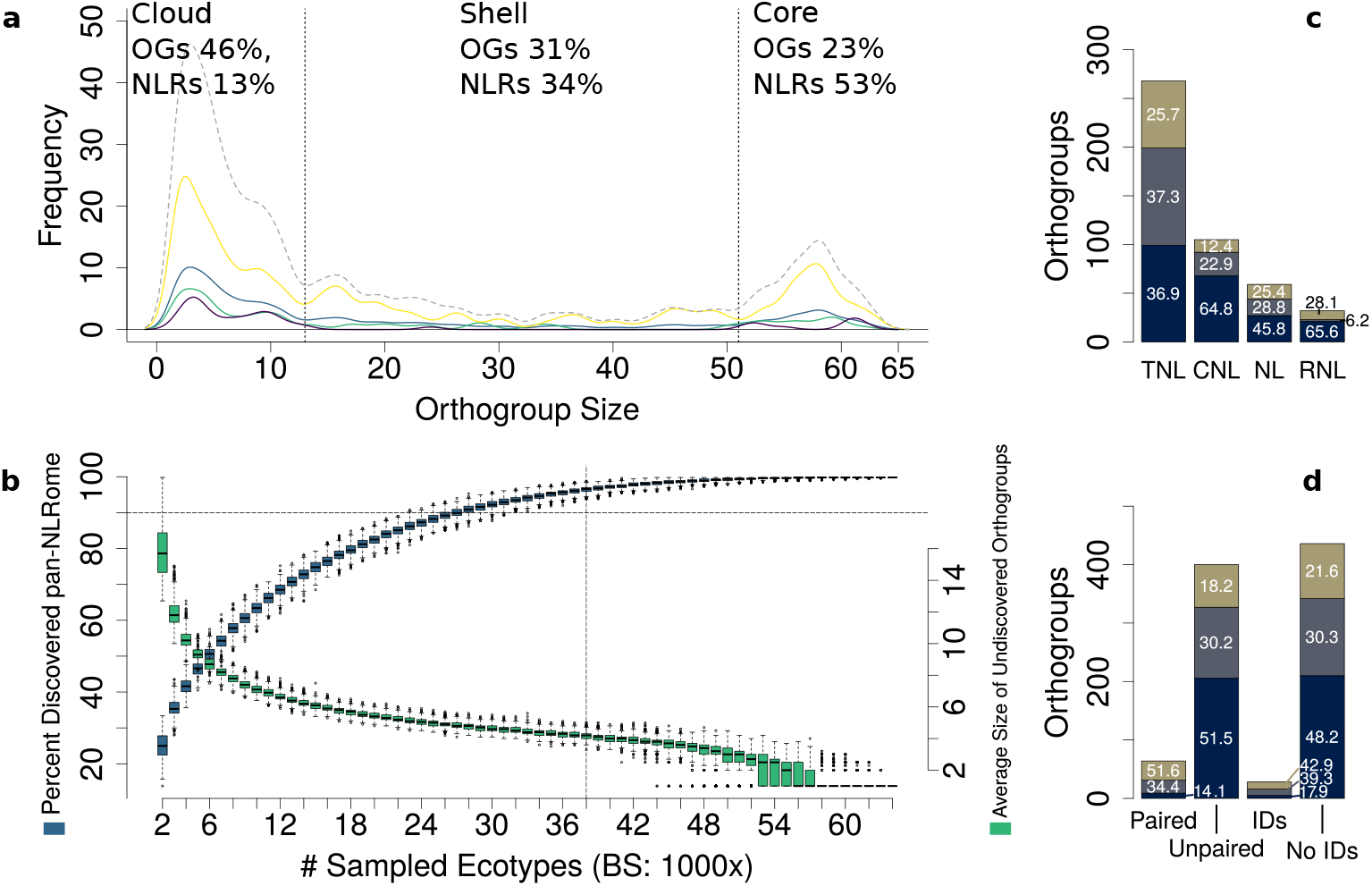
OG sizes, saturation, and distribution of core-, shell- and cloud-NLRs. a) OG size distribution for TNLs (yellow), NLs (green), CNLs (blue), RNLs (purple), and all NLRs (grey dashed line). The vertical lines at x=13 and x=51 differentiate cloud, shell and core. b) Saturation of the pan-NLRome. The blue boxes show the percentage of the pan-NLRome that can be recovered when randomly drawing a fixed number of accessions (1000x bootstrapping). The horizontal dashed line indicates where 90% of the pan-NLRome is found. The green boxes shown for each subset of drawn accessions indicate the average size of undiscovered orthogroups. The vertical dashed line indicates that 95% of the pan-NLRome can be recovered with 38 accessions. c) OG-type specific distribution of NLR classes in the cloud (dark blue), the shell (grey), and the core (olive green), and the relative fraction (numbers in the bars). d) OG-type specific distribution of paired and unpaired NLRs, and NLRs with and without IDs in the cloud (dark blue), the shell (grey), and the core (olive green), and the percentage (numbers in the bars).

### Placement of non-reference OGs

We discovered 296 high confidence OGs without a reference Col-0 allele, with six belonging to the core, 205 to the cloud, and 85 to the shell NLRome. In order to anchor these OGs to the reference genome, we asked how often orthogroups co-occurred, using OGs with known location (NLR and non-NLR OGs with a Col-0 reference allele) to anchor contigs with OGs lacking a reference allele. Anchoring efficiency of newly discovered OGs was calculated for co-occurrences in 2 to 39 accessions (Supplementary Table 10). With a minimum threshold of 10 accessions, we derived 42 co-occurrence subnetworks (Supplementary Fig. 4), anchoring 24 out of 132 OGs present in at least 10 accessions, but missing from the Col-0 reference. Most were anchored to other NLRs (Supplementary Table 10 and Supplementary Fig. 4). Newly anchored OGs include one CNL pair and three TNL pairs (Fig. 4, Supplementary Fig. 4, and Supplementary Fig. 5), with one ID-containing sensor-type OG (205.1) arranged in head-to-head orientation to the executor-type OG 204.1 (Supplementary Fig. 5). The use of annotated non-NLR genes in the assembled contigs allowed us to properly place these novel OGs.

**Figure 4.**
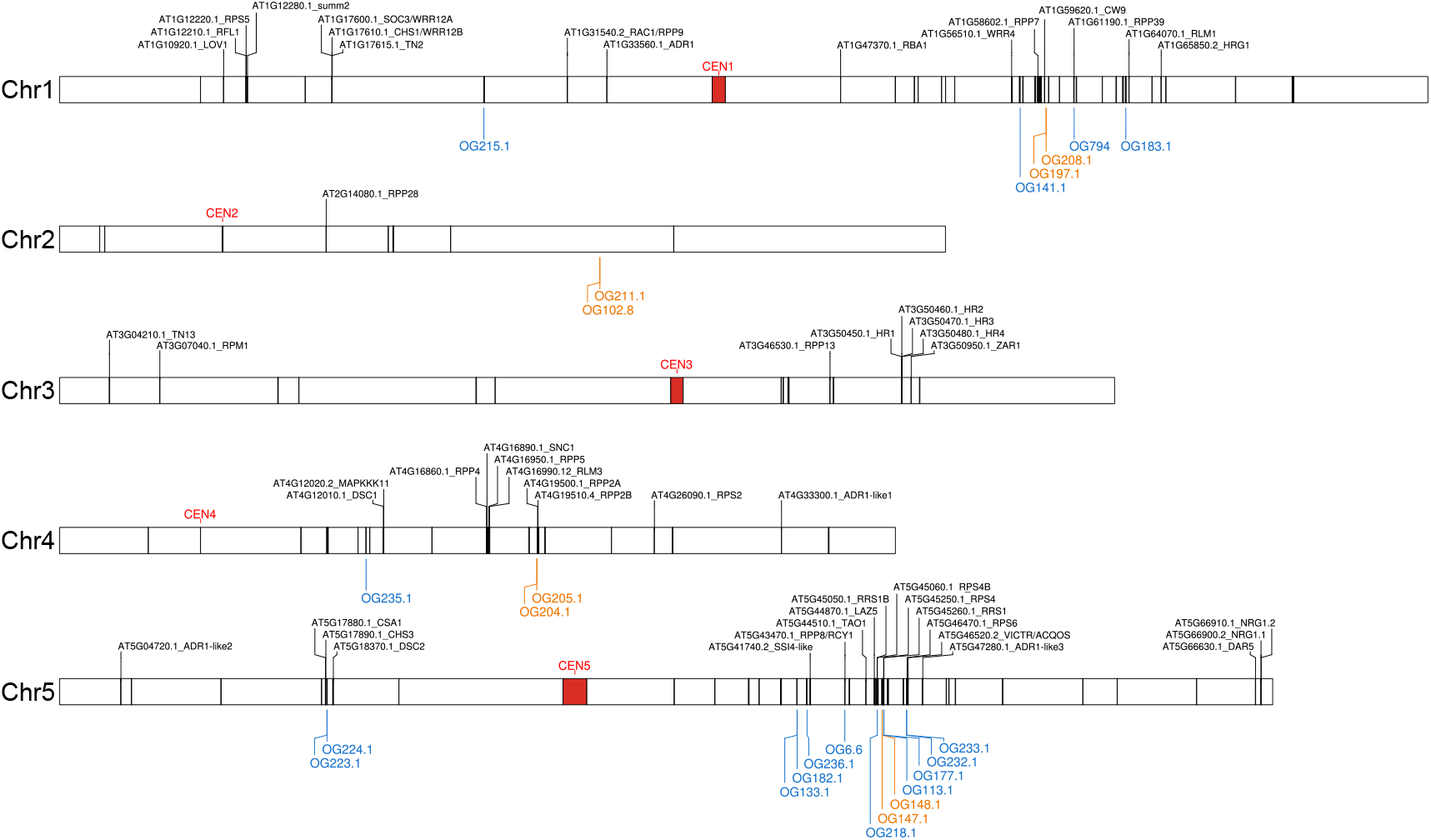
Genomic location of *NLR* genes in the reference assembly. The five *A. thaliana* chromosomes are shown as horizontal bars with centromeres in red. Reference NLRs are shown as black line segments. Text labels are shown only for functionally defined Col-0 NLRs. Anchored OGs found in at least 10 accessions are shown below each chromosome. Anchored OGs with paired NLR members are shown in orange, while remaining anchored OGs are shown in blue.

### Pan-NLRome Diversity

In an orthogonal approach to classifying NLR genes by their architectures, we assessed sequence diversity, an indication of the evolutionary forces shaping the pan-NLRome. Average nucleotide diversity was similar for CNLs, NLs and TNLs (Fig. 5a). It was lowest in RNLs, but because this is the smallest group, this difference is not statistically significant. The same trend was true for haplotype diversity (Fig. 5c). Nucleotide diversity was lowest in core and higher in shell and cloud OGs across TNLs, CNLs and NLs (Fig. 5a), suggesting that selection is relaxed in OGs with larger presence/absence variation. The pattern was reversed for haplotype diversity (Fig. 5c) and core OGs generally showed high values. We noticed that a few shell TNL and NL OGs stand out because of their ultra-low haplotype diversity, suggesting a conserved but rarely encountered selective pressure (without any correlation between geographic location and the accessions carrying these orthogroups). The average nucleotide diversity saturated with 32 randomly selected accessions, and the haplotype diversity with 49 accessions, suggesting a prevalence of low frequency haplotypes (Supplementary Fig. 6). Compared to non-clustered OGs, physically clustered OGs had significantly higher nucleotide diversity (Supplementary Fig. 3). This finding may indicate relaxed selection after gene duplication in these clusters^55^. When considering different NLR protein domains, the highest diversity was found in LRRs across all major classes and subclasses (Fig. 5b). Combining population genetics statistics for a Principal Component Analysis (PCA) revealed that more than 60% of the variance can be explained by as little as two components (Supplementary Fig. 7). However, none of the collected metadata, such as orthogroup size, type, class or the presence of IDs or a potential partner, explained the clustering of the first two principal components (Supplementary Fig. 7). This suggests a complex interplay of the different factors driving NLR evolution. Tajima’s D values, which can indicate balancing and purifying selection^56^, were similarly distributed across different NLR classes, with all classes containing extremes in both directions (Fig. 5d). Low Tajima’s D values were most common in TNLs, largely driven by core-and shell-type OGs.

**Figure 5.**
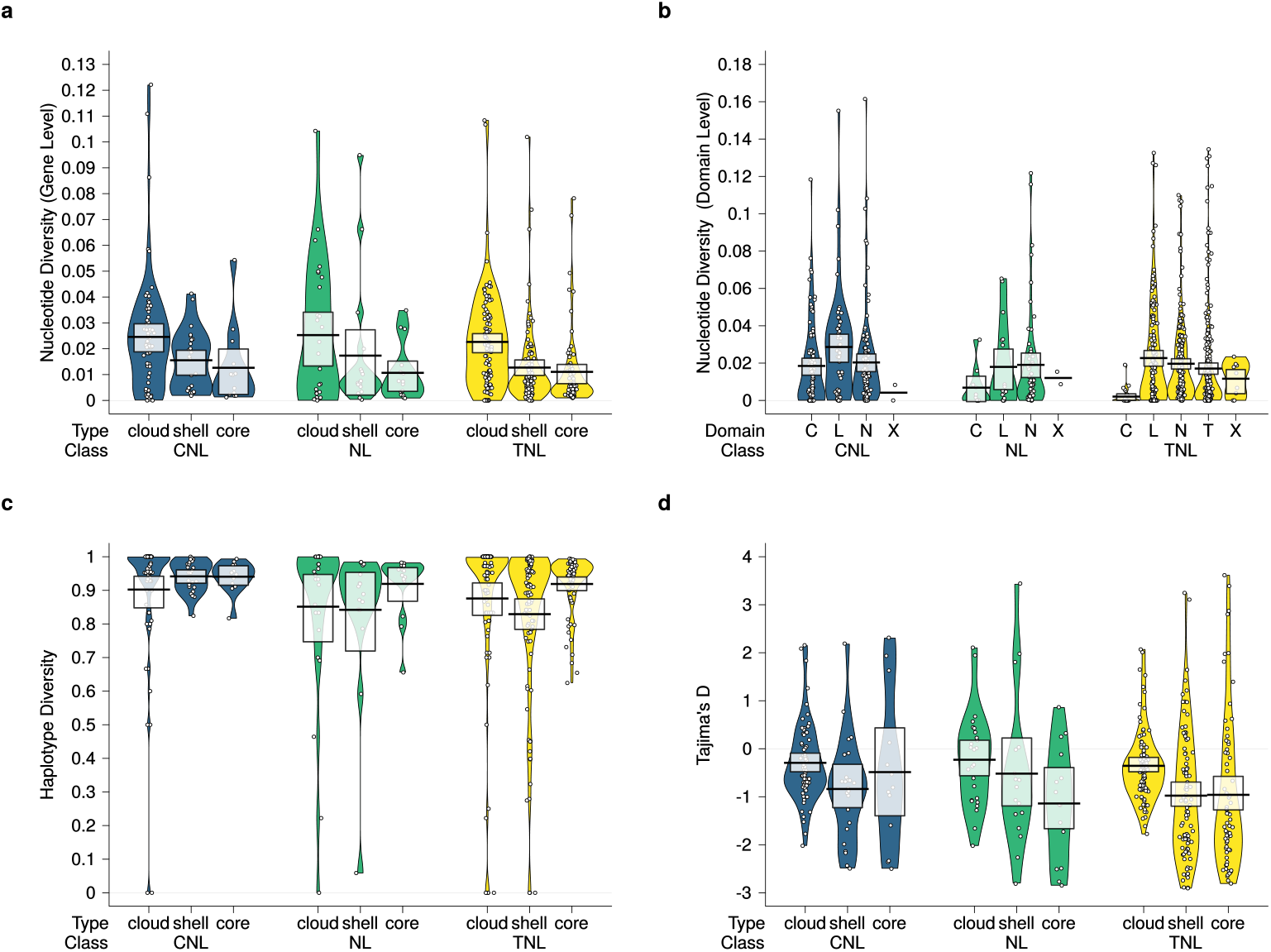
Population genetic statistics across the Pan-NLRome. a) Nucleotide diversity distribution grouped by orthogroup type and NLR class. Nucleotide diversity was defined as average pairwise nucleotide difference normalized to the number of sites in the respective orthogroup. b) Nucleotide diversity distribution grouped by domain type and NLR class. Sites of each orthogroup annotated with the same domain were aggregated. Type C within class NL occurs where a minority of OG members had an identifiable CC-domain. c) Haplotype diversity distribution grouped by orthogroup type and NLR class. Haplotype diversity was defined as average pairwise haplotype differences. A value of 1 indicates a high chance of finding two different haplotypes for two randomly chosen sequences of a given orthogroup. d) Tajima’s D (a measure of genetic selection) distribution grouped by orthogroup type and NLR class. RNL orthogroups are not shown because of the low number of orthogroups showing this class.

Site-specific selection analyses revealed core and shell OGs that have likely experienced constant (248), pervasive (165) or episodic (130) positive selection (Fig. 6a, b; Supplementary Fig. 8). Codons completely invariable, indicating constant positive selection, can be found across all types (e.g., core, shell) and classes (e.g., TNLs, CNLs). Pervasive positive selection seemed more likely in core-like OGs (71%) than in shell-like OGs (63%) while episodic positive selection patterns showed at a similar rate (52%). Subclasses showed a more uneven pattern of positive selection (Fig. 6e). Sites under constant positive selection were mostly found in NB, TIR and LRR regions when comparing all annotated protein domains (Fig. 6f). Pervasive and episodic positive selection patterns appeared predominantly in NB and TIR domains (Fig. 6g, h). A few OGs stood out because of the large fraction of codons of annotated protein domains under positive selection, including *RPP13* which confers race-specific downy mildew resistance^57^ (Supplementary Fig. 8). Sites under positive selection were also found in 11 IDs, including WRKY, TCP, B3 and DA1-like domains (Fig. 6c). Notably, invariant sites were detected in the WRKY domains of all three OGs containing a WRKY, and in a surprisingly high proportion of sites in the BRX domains of the RLM3-containing OG (Supplementary Table 11). We conclude that positive selection is widespread in the core-NLRome, being most prevalent in canonical NLR domains.

**Figure 6.**
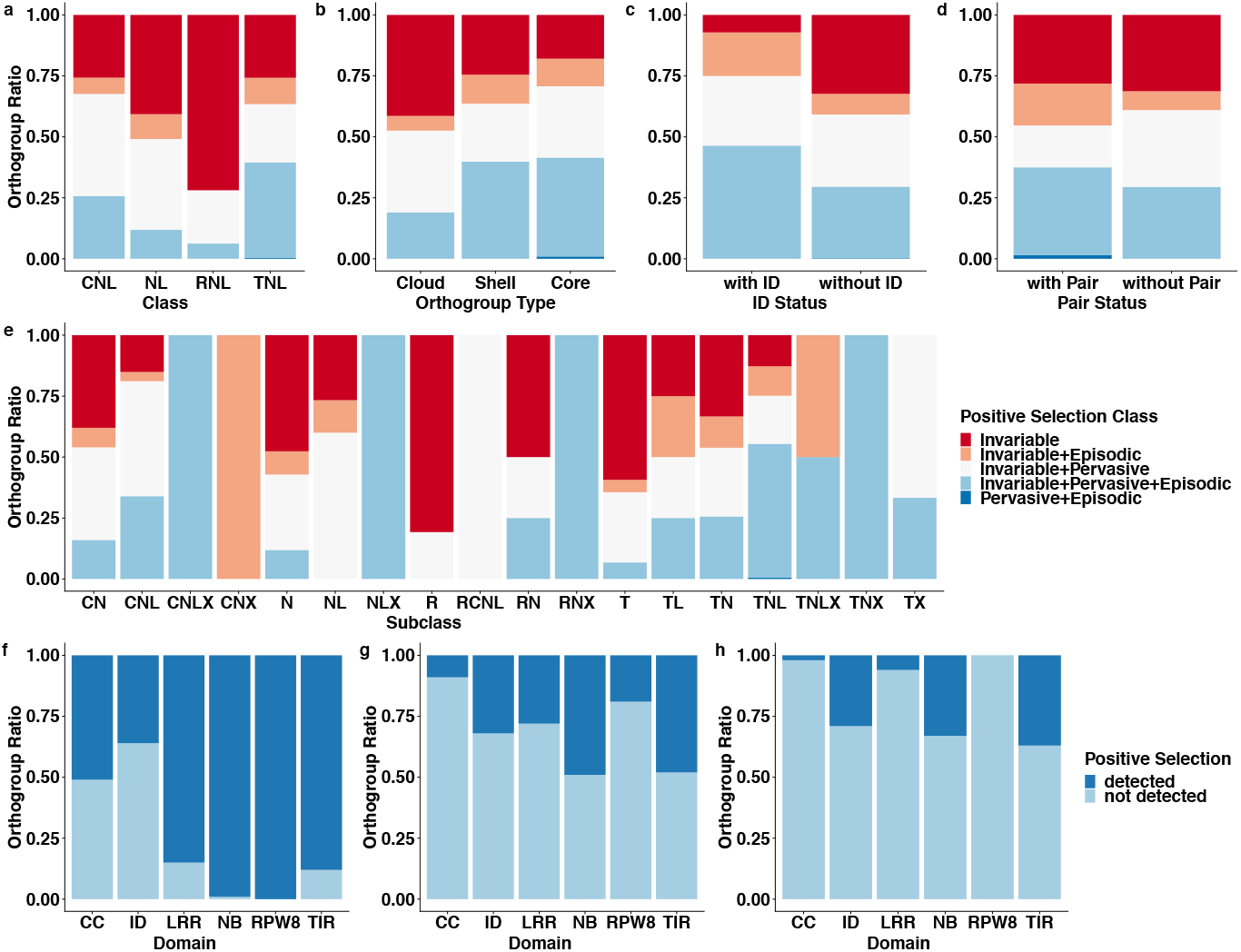
Positive selection landscape of the Pan-NLRome. (a-e) Ratio of different positive selection classes grouped by NLR class (a), orthogroup type (b), presence of a non-canonical domain (c), presence of a paired NLR (d) or NLR subclasses (e). An orthogroup was considered if at least one positive selected site of a given class was detectable. (f-h) Ratio between orthogroups showing constant (f), pervasive (g) or episodic (h) selection or no selection grouped by annotated protein domains.

### Linking Diversity to Function

Because NLRs that had been experimentally implicated in resistance to biotrophic pathogens showed enhanced diversity, we sorted OGs by resistance to adapted biotrophs (*Hyaloperonospora arabidopsidis*), non-adapted biotrophs (Brassica-infecting races of *Albugo candida*)^58^ and hemibiotrophs (mostly *Pseudomonas* spp.). OGs that provide resistance against adapted biotrophs are significantly more diverse than other categories (Fig. 7a; ANOVA and Tukey’s HSD p<0.01), suggesting that host-adapted biotrophic pathogens are driving diversification of NLRs more than other pathogens. That RNL helper NLRs have low diversity is consistent with their requirement to function with several sensor NLRs^59–61^.

**Figure 7.**
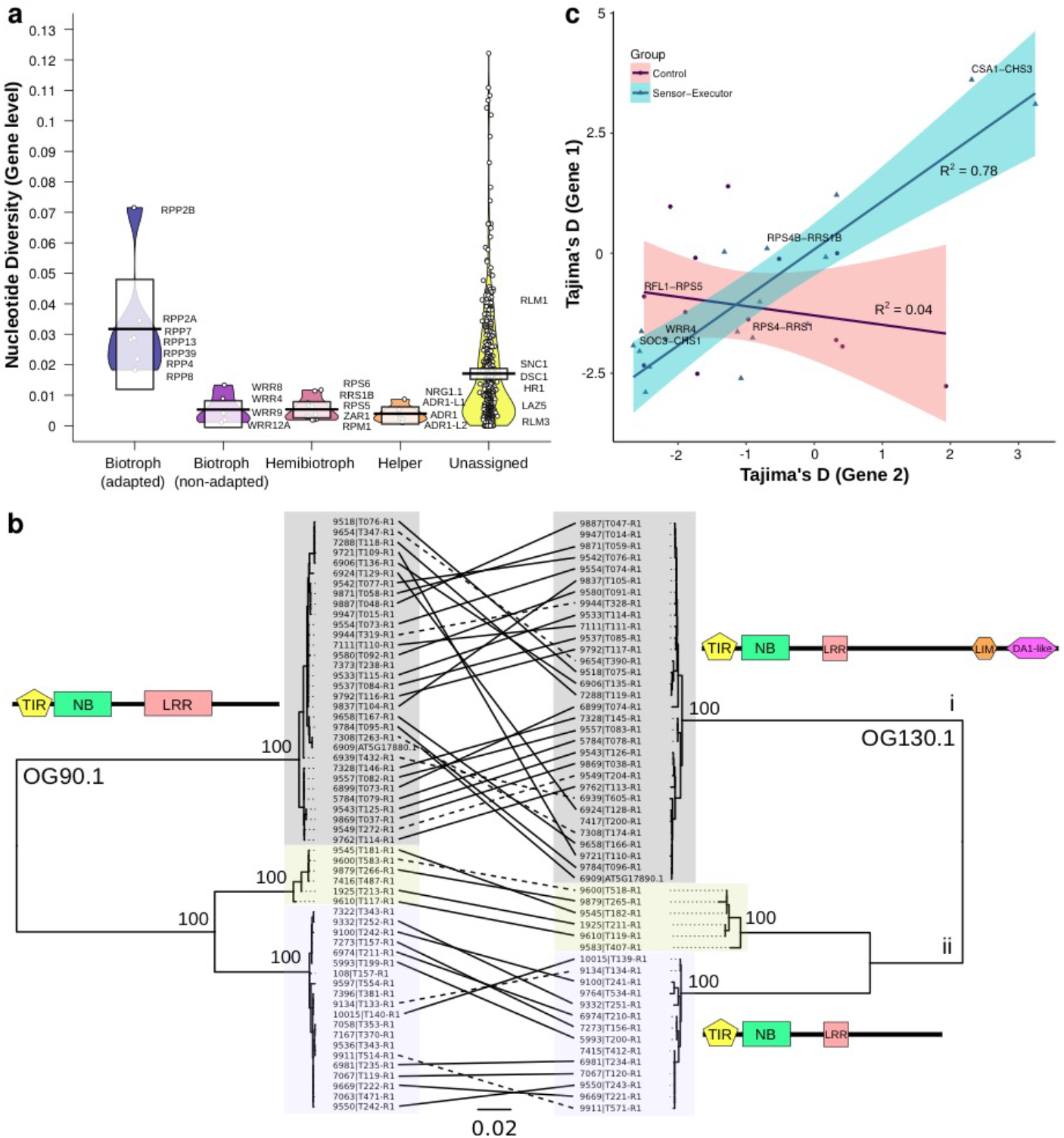
Population genetics of different OG classes grouped by known resistances and pairs. a) Nucleotide diversity distributions by functional class according to pathogen type to which they provide resistance. b) Correlation of Tajima’s D values in sensor/executor and control pairs. c) Maximum-likelihood phylogenetic trees of two OGs 90.1 and 130.1, which form a sensor/executor pair^63^ (100 bootstrap support indicated at major nodes). Scale bar refers to substitutions per site. Lines connecting the trees denote same accession.

Among the OGs with the lowest Tajima’s D values, a prominent example was *RPM1*, which confers resistance to a hemibiotrophic bacterial pathogen, and for which an ancient, stably balanced presence/absence polymorphism across *A. thaliana* is well established^62^. OGs that provide resistance to adapted biotrophs tend to have higher Tajima’s D values, indicating that they experience not only diversifying, but also balancing selection. Tajima’s D values within sensor-executor pairs encoded in head-to-head orientation were correlated whereas other closely linked NLR genes or random pairs were not (Fig. 7b, Supplementary Table 12). As an example, two OGs with high Tajima’s D values are the paired NLRs *CSA1* (OG91) and *CHS3* (OG130). *CHS3* featured two very different groups of alleles distinguished by the presence of LIM and DA1-like IDs^63^. This pattern was perfectly mirrored by the one for *CSA1*, the paired “executor” partner NLR of *CHS3* (Fig. 7c).

## Discussion

We defined the full species repertoire of the gene family that encodes NLR immune receptors in the model plant *A. thaliana*. Importantly, the pan-NLRome inventory became saturated with ~40 accessions randomly selected from the 65 accessions we analyzed. Before our work, it was known that there was excessive variation at some NLR loci, such that in the small number of accessions in which the relevant genomic region was analyzed in detail, every accession was very different, including significant presence/absence variation^41,64^. The fact that our pan-NLRome saturates with ~40 accessions indicates that the number of divergent loci is not unlimited. It also provides some guidance for future efforts in other species. It will be fascinating to compare the allelic and diversity saturation of self-fertilising *A. thaliana* with obligate out-crossers and with domesticated species. Among functionally annotated genes, we found the highest sequence diversity in NLR-coding genes whose products recognize evolutionarily adapted biotrophic pathogens.

We have also found an astonishing diversity of IDs, which allow hosts to rapidly accrue the ability to recognize the biochemical action of pathogen effector proteins. ID-containing NLRs that have been functionally characterized are all found in paired orientation. In these pairs, the ID member functions as pathogen sensor, and the other member as signaling executor^19–21,50,51,63,65^, with both members contributing to repression and activation of NLR signaling^66^. The correlation between Tajima’s D values of such paired NLRs support a co-evolutionary scenario whereby mutations into the sensor component lead to compensatory changes in the executor, or vice versa.

However, half of the 22 most commonly found IDs did not occur in an arrangement indicative of sensor/executor pairs. An open question is whether these function with unlinked executor partners, or whether they can function as dual sensor/executor proteins. Within the *A. thaliana* pan-NLRome, we identified three key families of defense-related TCP, WRKY and CBP60 transcription factors, represented as IDs in sensors of the class defined by RRS1. TCP domains are particularly interesting, as TCP transcription factors are preferentially targeted by pathogen effectors from divergently evolved pathogens^52,53,67,68^. The TCP domain may open a new avenue to engineering of NLR specificity, through TCP swap or inclusion of known effector-interacting platforms from TCP14^65^, as recently demonstrated with protease cleavage site swaps^30,67^. Furthermore, since many of the novel IDs were found at intermediate frequency in the population, and were novel compared to the Col-0 reference genome, we predict that this will apply to other plant species, suggesting that the number and diversity of NLR-IDs greatly exceeds that which has been so far reported.

## Supplementary Figure Legends

**Supplementary Figure 1. NLR frequency for different subclasses**. For each subclass, the corresponding class is color coded (TNLs: yellow, NLs: green, CNLs: blue, and RNLs: purple), and classes are in addition divided by the vertical dashed lines. NLRs are grouped into subclasses by their domains content: T (TIR), N (NB), C (CC), R (RPW8), and X (all other integrated domains). Each domain must be present at least once, domains in brackets may be present. Domain order is not considered. The mean is shown as a solid black horizontal line and the 95% Highest density Intervals (HDI: points in the interval have a higher probability than points outside) are shown as solid bands around the sample mean. All raw data points are plotted as open circles and the full densities are shown as a bean plot.

**Supplementary Figure 2. Schematic representation of NLR domain architecture diversity and simplification of consecutively repeated domains**. a) Examples of NLR domain architecture diversity. On top, a generic NLR, with an ID (Integrated Domain) is shown at the C-terminus. IDs can also be found at the N-terminus, and more rarely between the three canonical domain types. b) Reduction of domain combinations by collapsing duplicated/repetitive domains. The number of NLRs grouped by each of the original architectures is shown on the left, along with one example that can be visualized in the genome browser. Ellipsis in the bottom left represent 19 other architectures containing 4,079 proteins exclusively composed of TIR, NB and LRR domains. The same strategy was applied to all other architectures containing at least one duplicated domain in the RPW8, NB and CC classes. c) Full set of the novel *A. thaliana* NLR architectures. Includes the architectures contributed by only one gene. Domain architectures are shown in the y-axis. The number of NLRs in each architecture is shown in the x-axis. Asterisks indicate the 49 architectures not yet detected in the Brassicaceae outside of *A. thaliana*, or in the reference accession Col-0. Numbers next to y-axis show the ratio of paired NLRs divided by the total number of NLRs in each architecture.

**Supplementary Figure 3. OG size distribution comparisons**. Vertical black lines divide cloud (left section) from shell (middle section) and core (right section) NLRs. a) Comparison of OG size distributions of TNL OGs (blue) and CNL OGs (green) b) Comparison of paired (blue) and non-paired (green) OGs. c) Comparison of clustered (blue) and non-clustered (green) OGs. d) Comparison of ID-containing (blue) and non-ID-containing OGs (green). e) Distribution of Paired NLRs and NLRs with IDs across the Cloud- (dark blue), the Shell-(grey), and the Core- (olive green) pan-NLRome.

**Supplementary Figure 4. Orthogroup co-occurrence network**. Annotated NLR (green nodes) and non-NLR genes (white nodes) clustered into OGs were analyzed for co-occurrence in the same contig. The number of co-occurrences is represented by grey lines connecting nodes (edges). The minimal co-occurrence threshold imposed was 10 accessions, but similar networks can be derived for any number accessions. NLR OGs without a Col-0 allele (green square nodes) are highlighted in blue boxes. Hypothetically paired OGs not known in Col-0 are highlighted in orange boxes.

**Supplementary Figure 5. Quantitative co-occurrence of the novel hypothetical paired NLRs in OG205.1 and OG204.1**. Abbreviations: OG, Orthogroup; H2H, Head-to-head; NA, Not available.

**Supplementary Figure 6. Nucleotide and haplotype diversity saturation**. Saturation of nucleotide (a) and haplotype (b) diversity after random subsetting of the complete Pan-NLRome into bins of increasing sizes. For each size 100 bootstrap were carried out.

**Supplementary Figure 7. Population genetics statistics based PCA**. Principal component analysis carried out on 10 population genetics statistics, namely Nucleotide diversity / Pi, Haplotype diversity, Fu and Li’s D, Fu and Li’s F, Tajima’s D, Rozas’ R_2_, Strobeck’s S and the number of segregating sites. Panels are colored according to categorical variables.

**Supplementary Figure 8. Positive selection landscape of the Pan-NLRome**. (a-e) Absolute number of orthogroups in positive selection classes grouped by NLR class (a), orthogroup type (b), presence of a non-canonical domain (c), presence of a paired NLR (d) or NLR subclasses (e). An orthogroup was considered if at least one positive selected site of a given class was detectable. (f-i) Domain coverage with positively selected sites grouped by NLR class and positive selection type across canonical domains (f-g, i) and all aggregated non-canonical domains (h).

**Supplementary Figure 9. Read and assembly statistics**. a) Read lengths distribution (Q20-filtered CCS reads) for all accessions (black). The mean is shown as a solid black horizontal line. The full densities are shown as a bean plot. The total number of CCS reads (blue circles) and the total number of bases (orange diamonds) are plotted in addition. b) Contig lengths distribution (black). The mean is shown as a solid black horizontal line and the 95% Highest density Intervals (HDI: points in the interval have a higher probability than points outside) are shown as solid bands around the sample mean. The full densities are shown as a bean plot. Raw data points are plotted using black dots. The total assembly sizes (orange circles) are plotted in addition. c) Quality (black) and completeness values (orange) for sub-sampled Col-0 datasets. The amount of input data for each sub-sampling experiment is shown as a second x axis. d) Quality (black) and completeness values (orange) for all RenSeq accessions. Unfilled circles indicate accessions with qualities higher than any sub-sampled dataset. The vertical black line is drawn at 95% completeness. e) Correlations between the Assembly Quality, the amount of Input Reads, the amount of Input Bases [bp], the read length N50 [bp], and the similarity to Col-0 are shown for the RenSeq datasets. Histograms and kernel densities (red line) are plotted for each variable. Scatter plots for variable pairs are shown together with a fitted line (red) and the Pearson’s correlation coefficient (significance 0 ‘***’,0.001 ‘**’,1 ‘ ’).

**Supplementary Figure 10. Phylogenetic tree of NB domain alignments for refining sensor/executor pairs from all pairs**. RPS4/RRS1-like and SOC3/CHS1-like paired TNLs fall into distinct subclades. These are indicated by color: RPS4-like (Silver, executors), RRS1-like (Gold, sensors), SOC3-like (Pink, executors) and CHS1-like (Bronze, sensors). This phylogeny was constructed by aligning the NB domain (~240 amino acids) of all TIR and NB containing Col-0 proteins and selected additional representatives of pair flagged orthogroups (OGs) from the pan-NLRome that are not represented in Col-0 (identified by their OG and protein numbers). NB domains from APAF1 (Human) and AT1G58602.1 (A. thaliana CNL) were also included. Amino acid sequences were aligned with MUSCLE (Neighbor joining clustering), refined by manual trimming and the phylogeny produced with the WAG maximum likelihood method allowing for 3 discrete Gamma categories. AT4G36140 contains two distinct NB domains, both of which were included and the second of which clusters with other RRS1-like NB domains. Number of 100 bootstraps supporting topology shown at major node vertices. Scale bar represents amino acid substitutions per site.

**Supplementary Figure 11. Orthogroup (OG) size frequencies before OG refinement**. Data are shown separately for the different NLR classes (TNL: yellow, NL: green, CNL: blue, RNL: purple) and for OGs with (solid lines) and without (dashed lines) complex paralogs (duplications spread across the whole phylogeny).

## Supplementary Tables

Supplementary Table 1. 65 accessions metadata. Column content explained in the github repository.

Supplementary Table 2. Domains found in the *A. thaliana* pan-NLRome. Column content explained in the github repository.

Supplementary Table 3. Gene models included in the *A. thaliana* pan-NLRome. Column content explained in the github repository.

Supplementary Table 4. Universe of IDs detected in the *A. thaliana* reference Col-0 accession and/or a 19 other Brassicaceae NLRomes.

Supplementary Table 5. Gene models used to generate the Brassicaceae NLRome.

Supplementary Table 6 Sources of the 22 proteomes from 19 Brassicaceae species used to generate the Brassicaceae NLRome. see Online Methods.

Supplementary Table 7. Domain architecture prevalence, number of paired NLRs and presence in the *A. thaliana* reference Col-0 accession and/or 19 other Brassicaceae NLRomes.

Supplementary Table 8. Number of architectures detected for each of the 27 IDs already known from 22 Brassicaceae NLRomes.

Supplementary Table 9. Architecture metadata. Columns are explained in the github repository.

Supplementary Table 10. Number of non-reference OGs that can be anchored to Col-0 genomic positions through same contig co-occurrence with reference OGs.

Supplementary Table 11. Frequency of positively selected sites in putative integrated domains.

Supplementary Table 12. List of orthogroups categorized as sensor/executor or control pairs, with functional metadata and Tajima’s D.

Supplementary Table 13. Oligos used to introduce custom barcoded adapters in TSL accessions.

Supplementary Table 14. Bait library (v2.4) used to capture NLR-coding genes.

Supplementary Table 15. First and second subdivisions used to define NLR classes.

Supplementary Table 16. Prediction of coiled-oil motifs in functionally validated Col-0 CNLs.

Supplementary Table 17. Homology-based clustering into orthogroups without refinement

Supplementary Table 18. Pervasive diversifying positive selection posterior probabilities

Supplementary Table 19. Positional pervasive diversifying positive selection posterior probabilities

Supplementary Table 20. Episodic diversifying positive selection p-values

Supplementary Table 21. Positional episodic diversifying positive selection p-values

Supplementary Table 22. Refined final Orthogroups

Supplementary Table 23. Enrichment of curation flags in orthogroups calculated. Only q-values below 0.1 are reported.

Supplementary Table 24. Misannotated genes manually removed from the NLRome.

Supplementary Table 25. Non-NLR Orthogroups used for non-reference OG placement.

## Data Availability Statement

Raw data and assembled sequences were deposited at the European Nucleotide Archive (ENA) under accession number PRJEB23122. Genome browser is available at http://ann-nblrrome.tuebingen.mpg.de/annotator/index. Manually curated gene models(gff), domain annotations, orthogroups, protein and transcript alignments, phylogenetic trees, scripts necessary to produce figures and further metadata files containing information parsed and restructured from the supplemental tables in this manuscript are available at github (https://github.com/weigelworld/pan-nlrome/). Visualization of OG phylogenetic trees and metadata is available at iTOL (https://itol.embl.de/shared/pan_NLRome).

## Acknowledgements

We thank Florian Jupe for contributing with methods before publication. Eunyoung Chae for contributing with alleles for bait design. Burkhard Steuernagel for assistance with demultiplexing. Johannes Hofberger and Eric Schranz for providing sequences used in a former version of the bait library. JLD is an Investigator of the Howard Hughes Medical Institute (HHMI). This work was supported by a grant from the Gordon and Betty Moore Foundation to the 2 Blades Foundation (GBMF4725), and the Howard Hughes Medical Institute (HHMI). OF, JJ and KW acknowledge support from the Gatsby Charitable Foundation.

## Glossary

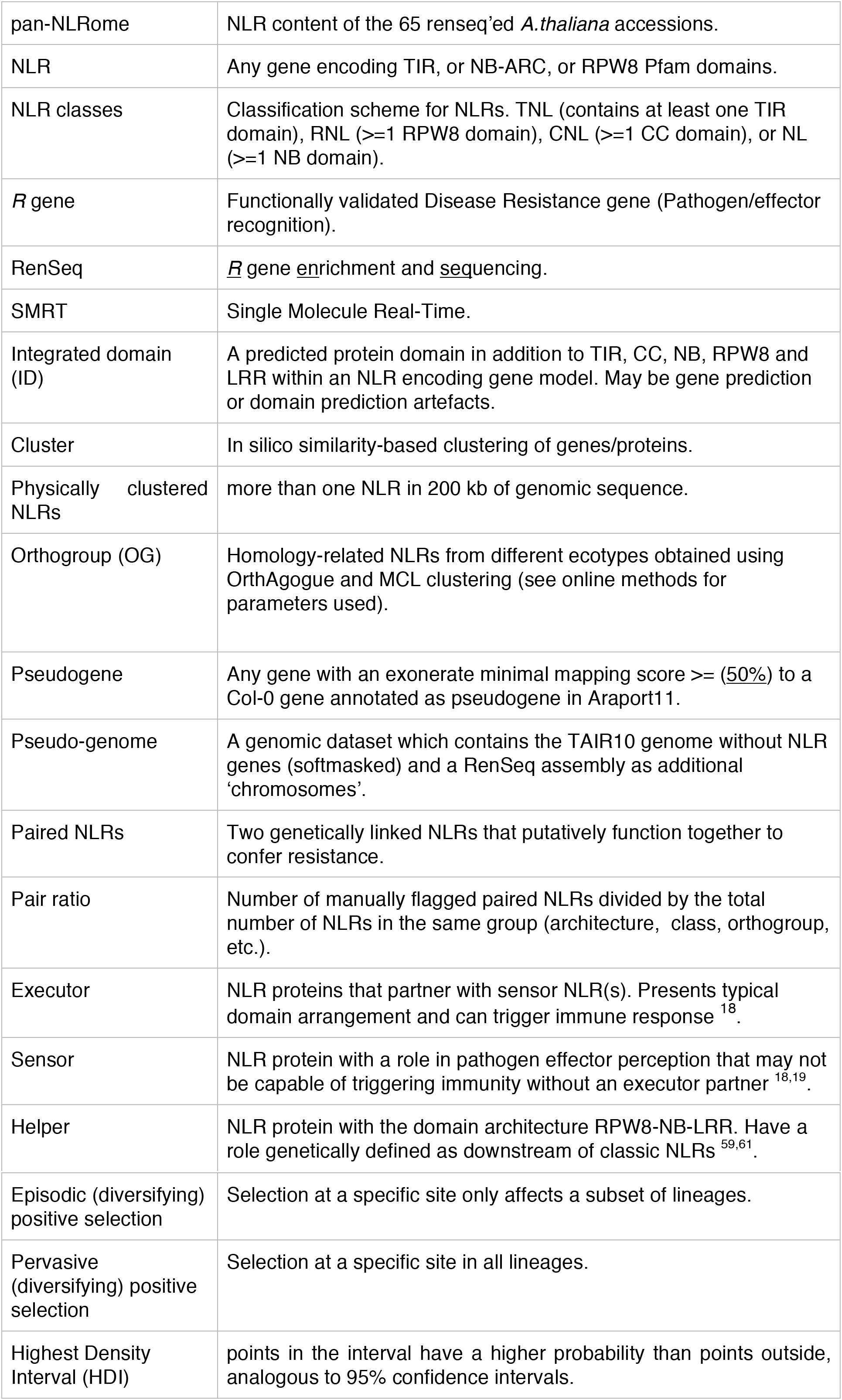

## Author Contributions

Project Conception JJ, JLD, DW

Project Management FB, FM, AVDW, OF

Bait Design OF, MN, VC

Pre-publication methods access KW

Data Generation AVDW, FM, MN, OF, VC

Data Preparation & Assembly AVDW, FB, FM

Gene Annotation AVDW, FB

Gene Curation AVDW, FM, OF

Architecture Analysis FM

Genomic Placement FM

OG Co-occurrence network FM

Pan-NLRome Generation FB

Population Genetics Analysis FB, OF

Positive Selection Analysis FB

Metadata Collection FB, AVDW, DW

Data Visualization (iTOL, JBrowse) FB, AVDW

Initial draft manuscript FB, AVDW, FM, OF

Manuscript Revision FB, JLD, OF, JJ, FM, AVDW, DW, MN, VC

