## Supplementary material for "The *Arabidopsis thaliana* pan-NLRome": Online Methods and Supplementary Figures

Supplementary figures may be found at the end of this document.

### Online Methods

We characterized NLR gene variation in *A. thaliana*. NLRomes were generated for a diverse set of 65 ecotypes by targeted sequencing of long NLR containing genomic fragments. Resistance gene enrichment sequencing (RenSeq) is a targeted enrichment strategy that uses synthetic biotinylated RNA probes to capture DNA based on similarity [1]. Witek and collaborators combined RenSeq with PacBio (SMRT RenSeq), to obtain long and curated reads that were used to unambiguously define NLR clusters, map and clone novel resistance genes [2]. This method allows the specific sequencing and assembly of an organism's NLR gene complement, and several kilobases of flanking DNA sequences.

#### *De novo* assembly and NLR annotation

##### Ecotype Selection

*Arabidopsis thaliana* accessions were selected to attempt to cover the species' NLR gene diversity. For that, we sequenced the NLRome of 65 accessions that include a subset of 20 naturally occurring diverse accessions known as 'relicts' (mostly Iberian accessions that contain an unusually high amount of genetic diversity) [3], a subset of 19 in-lab selected phenotypically diverse panel known as MAGIC founders [4,5], and a set of remaining accessions with high diversity on whole genome level and representing the different known haplotypes [3] (Supplementary Table 1).

##### Ecotype Verification

Routine seed stock genotyping prevents sample contamination [6]. At a late stage of this project, 46 accessions were re-sequenced as part of a routine seed stock verification effort and the ecotypes were determined using SNPmatch as described [6]. Three mis-labelled accessions were found in our dataset (Identifier: 7063, 9911 and 9658). Their ecotype names and ecotype IDs from the 1001 genomes project were corrected and reported (Supplementary Table 1). For the sake of contiguity, their identifiers were not changed.

##### SMRT RenSeq

Genomic libraries were enriched for NLRs and sequenced using PacBio long read technology [7]. Library preparations were performed collaboratively in three labs (UNC, MPI, and TSL) with minor handling differences. DNA was extracted and fragmented to 2-5kb pieces for long read circular consensus sequencing (CCS). It was prepared using either the DNeasy plant Maxi kit (UNC) (Qiagen, CA, USA), a custom high molecular weight DNA extraction protocol (MPI), or grinding in Shorty buffer (20% 1M Tris HCl pH 9, 20% 2M LiCl, 5% 0.5M EDTA, 10% SDS, 45% dH2O), followed by phenol chloroform extraction and precipitation with isopropanol (TSL). DNA was fragmented into shorter pieces using either Covaris red miniTubes (Intensity=1, DutyCycle=20%, Cycles per Burst=1000, Treatment time=600s, Temperature=20°C, Water level=15, Sample volume=200ul) (TSL), or Covaris g-tubes using manufacturer instructions for a targeted size of 6 kb (UNC, MPI) (Covaris, MA, USA). The DNA was purified using 0.4x AMPure XP beads (Beckman Coulter, IN, USA) according to the manufacturer's instructions.

Libraries for NLR enrichment were constructed using 'NEBNext Ultra DNA Library Prep Kit for Illumina' (New England Biolabs Inc, MA, USA). Sixteen accessions from TSL were prepared for multiplexed sequencing, by introducing custom barcoded adapters (dual 8 bp index) instead of the standard ones (Supplementary Table 13). For the PCR amplification, 5-10 $\mu$l adaptor-ligated DNA was used together with 25 $\mu$l 2x KAPA HiFi HotStart ReadyMix, 1 $\mu$l Index and Universal PCR Primer, and 13-18 $\mu$l water (to a total volume of 50 $\mu$l) (Kapa Biosystems, MA, USA). Initial Denaturation (94 °C for 4 min) was followed by at least 8 cycles (denaturation: 94 °C for 30 sec, annealing: 65 °C for 30 sec, extension: 68 °C for 4 min) and a final extension (68 °C for 10 min).

The genomic libraries were enriched for NLR genes. Roughly, 1.4 Mb of the reference Col-0 *A. thaliana* genome encodes NLRs. Baits were designed to hybridise with NLR-coding genomic DNA regions, and only bound fragments were sequenced (Supplementary Table 14). 20,000 synthetic 120 nt biotinylated RNA probes (*bait library*), complementary to 736 known NLR genes from *Arabidopsis thaliana* TAIR10 [8], *Arabidopsis lyrata* [9], *Brassica rapa* [10], *Aethionema arabicum* [11] and *Eutrema parvulum* [12] were ordered as a MYbaits kit (MYcroarray, MI, USA) (Supplementary Table 14). For manually selected Arabidopsis genes additional alleles were included, along with non-repetitive intron regions to improve capture in genes with introns longer than 350 bp. 100-500 ng of the libraries were hybridized with the baits using half of the reaction volume suggested in MYbaits v3.0 protocol with the following modifications: For each capture reaction, hybridization mix was prepared using 10 $\mu$l Hyb#1, 4 $\mu$l Hyb#3, 0.4 $\mu$l Hyb#2 and 0.4 $\mu$l Hyb#4; library mix with 2.5 $\mu$l SeqCAP (Roche), 0.3 $\mu$l Block#3 and 3 $\mu$l gDNA Library; capture mix with 2.5 $\mu$l Bait library and 0.5 $\mu$l RNase block (MYcroarray, MI, USA). Following the manufacturer's cycling conditions, we brought the mixes to a hybridization temperature of 65 °C and transferred 5 $\mu$l of the library mix and 5.5 $\mu$l of the hybridization mix to the baits. After 16 to 24 hours hybridization the enriched libraries were recovered using 50 $\mu$l Dynabeads MyOne Streptavidin C1 beads (Life Technologies, CA, USA). Binding and washing was carried out according to Mybaits 3.0 manual without the use of Hyb#4. Incubation of the captured libraries with the streptavidin beads was increased to 45 minutes. 30 $\mu$l molecular biology grade water was used to re-suspend the DNA. The captured libraries were amplified for 18-30 cycles using the KAPA HiFi DNA Polymerase and the protocol for cycling conditions given in the previous paragraph (Kapa Biosystems, MA, USA).

Libraries were prepared for long read sequencing. PacBio libraries for MPI data were prepared using the '2 kb Template Preparation and Sequencing' protocol (Pacific Biosciences, CA, USA), and were size selected for 2-5 kb using a BluePippin (0.75% Agarose Dye-Free/0.75% DF 2-6kb Marker S1, Start=2000, End=2000) (Sage Science, MA, USA). PacBio library prep for UNC data was done using the manufacturer's recommended procedure for '5 kb Template Preparation and Sequencing', and size selection for fragments over 3kb was done using a SAGE-ELF apparatus using 0.75% gel cassettes (Sage Science, MA, USA), size-based separation mode, target value 3 kb and target well 10. All wells containing fractions above 3 kb were pooled. Libraries for TSL data were prepared by size selecting fragments >3 Kb from the captured library using a SAGE-ELF apparatus as described above.

Quality control of all libraries was performed using the Qubit quantitation platform (Life Technologies, CA, USA) and Bioanalyzer (Agilent, CA, USA). The PacBio RS II sequencing platform and P6-C4 chemistry was used to sequence each accession or multiplexed pool on individual SMRT cells (Pacific Biosciences, CA, USA). Sequencing of several accessions was repeated in order to obtain sufficient output reads (Supplementary Table 1).

#### Read Correction

Raw reads were used to produce highly accurate corrected reads. Circular consensus sequencing produced overlapping raw reads that were self-corrected to consensus reads which reduces the read error from 17% to 2% (CCS; version 2.0.0; defaults [13]). Where indexing was employed, corrected sequences were de-multiplexed using a custom script available on github (see demultiplexing script at https://github.com/weigelworld/pan-nlrome/). One combined CCS read dataset was created for accessions that were sequenced on more than one SMRT cell. Only CCS reads with more than 99% per base accuracy were considered further (Supplementary Fig. 9a).

#### Assembly

Reads were assembled to rebuild NLR containing regions (Canu; version 1.3; -pacbio-corrected, trimReadsCoverage=2, errorRate=0.01, genomeSize=2m [14]). Expected genome size was adjusted to 2Mb which reflects the proportion of the genome captured with RenSeq according to the reference genome TAIR10 (1.4Mb NLR genes + expected flanking regions). Error rate reflected the input data quality after read correction. Read ends were trimmed using a minimum evidence of two reads. Contigs were removed if they were fully contained in a larger contig with >99.5% identity. The final assembly size and contig length distribution can be seen in Supplementary Fig. 9b.

#### Assembly Validation

##### Quality Scores

We used pseudo-heterozygous SNP calls created by mis-mapped reads, as a measurement for assembly quality. A read that couldn't be mapped to its correct NLR origin because the NLR was not assembled, instead mapped to a similar NLR and created pseudo-heterozygous SNPs. The quality was calculated from the ratio of pseudo-heterozygous SNPs and the total amount of mapped bases.

The pseudo-heterozygosity was determined using the corrected CCS reads. A pseudo-genome was constructed for each RenSeq assembly by combining the assembled contigs with chromosomes from the tair10 reference. To avoid mis-mappings, non-NLRs were masked on the RenSeq contigs, and NLRs were masked on the reference chromosomes. CCS reads were then mapped to those pseudo-genomes (minimap2; 2.9-r748-dirty; -x map-pb [15]). SNPs were called for NLR genes using high quality mappings only (htsbox pileup; r345; -S250 -q20 -Q3 -s5 [16]).

The number of pseudo-heterozygous sites (hetsites) was compared to the total number of mappable NLR gene bases (totalsites). The quality was calculated as logarithmically linked to the ratio of pseudo-heterozygous calls to the total amount of mapped bases (Supplementary Fig. 10d).

$$Q = abs(-10 * log10(\frac{hetsites}{totalsites}))$$

##### Completeness Assessment

The completeness of an accession's NLR complement was derived from quality and completeness relationships in the reference Col-0. We created the correlation between completeness and quality for Col-0 sub-assemblies using different amounts of input data. The corrected CCS reads from Col-0 were sub-sampled from 100% to 1% in 1% steps (seqtk sample; v.1.0-r82-dirty; defaults [17]). 100% of the data correspond to 26.639 reads with a N50 read length of 2.846 bases and 77.98 megabases sequence in total. The sub-sampled datasets were assembled with Canu (see pan-NLRome Generation for settings). All genes from the original RenSeq Col-0 assembly were mapped to each sub-assembly to detect assembled NLRs. NLR transcripts were extracted using those alignments (exonerate; v.2.2.0; --model est2genome --bestn 1 --refine region --maxintron 546 [18]). The quality of each sub-assembly was assessed as described above (Supplementary Fig. 10c). The completeness of a sub-assembly was determined as the fraction of the full reference Col-0 NLR complement that was assembled. NLR transcripts were evaluated using rnaQUAST (version 1.5.0; defaults [19]) with the TAIR10 reference genome and Araport11 NLRs. The completeness was calculated by dividing the amount of covered NLR genes (in bases) by the total length of the Araport11 NLRs (Supplementary Fig. 10c). The relation between completeness and quality of the tested Col-0 sub-assemblies was used to infer completeness values for the other accessions. Each accession's quality was used to find the corresponding completeness value from the tested Col-0 sub-assemblies (Supplementary Fig. 10d).

##### Similarity to Col-0

We determined if the similarity of an accession to the reference Col-0 influenced its quality (Supplementary Fig. 10e). RenSeq assemblies were mapped against the Col-0 assembly (minimap2; 2.9-r748-dirty; defaults [15]) and SNPs were called in NLR gene regions (htsbox pileup; r345; defaults [16]). Only biallelic SNPs were used to calculate the Identity By State (IBS) value for each accession compared to Col-0 (SNPRelate_1.10.2; method='biallelic' [20]).

#### Annotation

Coding and non-coding elements were annotated. Evidence and profile based methods were integrated in the MAKER pipeline (version 2.32; pred_flank=150, keep_preds=1, split_hit=3200, ep_score_limit=95, en_score_limit=95; [21]). Genes were predicted with AUGUSTUS (version 3.1.0; defaults [22]) and SNAP (version 2006-07-28; defaults; [23]). AUGUSTUS used the default 'arabidopsis' profile for gene prediction, and SNAP used a custom Hidden Markov Model (hmm) based on NB-ARC and/or TIR containing genes. Gene predictions were improved using Col-0 proteins and transcripts from the Araport11 website (Araport11_genes.20151202.pep.fasta, Araport11_genes.20151202.mRNA.fasta [24]).

Protein and transcript evidence was considered only if its mapping quality was high enough (see above for ep_score_limit and en_score_limit). Repeat-masked regions were not used for gene prediction (RepeatMasker; version open-4.0.5; model_org=arabidopsis; http://www.repeatmasker.org/RMDownload.html)

*Capsella rubella* and *Arabidopsis lyrata* reference annotations were revised to create reliable sets of NLRs for those outgroups. Reference annotations, evidence and gene predictions were integrated in MAKER. RNA-seq data guided gene prediction with BRAKER1 (version 1.9; defaults [25]). Reads from silique, root, stem, leaf, and flower (PRJNA336053; PE; 100bp; 5-10MB [26]) were mapped to the reference genomes using HISAT2 (version 2.0.5; --no-mixed --no-discordant [27]). Mapped reads guided gene prediction and were also used to assemble transcripts (Cufflinks; version 2.2.1; defaults [28]). Gene predictions were compared to reference gene annotations using MAKER (pred_gff, model_gff). Evidence mappings were used to choose the best annotation per locus. Reference genomes and annotations were taken from Phytozome (https://phytozome.jgi.doe.gov/ [29,30]). Assembled transcripts acted as the primary evidence (est_gff), re-annotated *A.thaliana* NLR transcripts and proteins were used as alternative evidence (altest, protein).

Protein domains indicate conserved functional, or structural, units of proteins. They were predicted for gene models and for AUGUSTUS gene-prediction products using Pfam hmms and coiled coils (InterProScan; version 5.20-59.0; -dp -iprlookup -appl Pfam,Coils [31]). Repeats often mark genomic regions with complicated annotations. RepeatMasker results were visualized. Diverged repeats in outgroups were additionally masked and visualized (repeat_protein=te_proteins.fasta provided by MAKER).

#### Web Apollo

Gene models and evidence tracks from Maker were integrated into WebApollo for manual inspection (version 2.0.4; http://ann-nblrrome.tuebingen.mpg.de [32]). Additional evidence tacks were added to evaluate the quality of the gene models. A track for duplicated and diversified genes was added by aligning transcripts (track=est2genome-50) and proteins (track=protein2genome-50) from the reference gene annotation (Araport11) to each NLRome (--percent 50, exonerate; version 2.2.0 [18]). The same procedure was carried out on known pseudogenes transcripts. Protein domain predictions were added for both Maker (track=InterProScan) and Augustus (track=InterProScan Augustus) gene models. A track with CCS read mappings (pbalign; version 3.0; defaults; [33]) was added to aid contig quality inspection. In case of *A. lyrata* and *C. rubella*, RNA-seq alignment data was added to inspect intron-exon boundaries.

#### Manual Re-annotation

Genes containing NB-ARC or TIR domains were manually inspected to create accurate and reliable annotations (see reannotation SOP at https://github.com/weigelworld/pan-nlrome). Gene models were evaluated using several biological evidence layers in Web Apollo. Incorrectly fused genes were split, and incorrectly split genes were merged. Col-0 protein and transcript mappings were used to detect wrongly fused or split gene models. Genes were split if several proteins or transcripts mapped next to each other within one model. Genes were merged if protein or transcript mappings spanned several models. Both cases often showed disagreeing gene predictions. Additional features of fused genes were extremely long introns, or pseudogene mappings. Evidence from protein and transcript mappings, as well as RNA-seq read mappings was considered to select the best gene model. Gene structures were corrected. Intron-, exon-, and UTR boundaries were refined. . Alternative splice forms were not used in this study. Genes were flagged with ‘corbound’ if exon-intron structures were changed without direct protein or transcript evidence, and ‘cortrans’ was used, if translation start points were changed (see gff files at https://github.com/weigelworld/pan-nlrome). Exceptions were detected and evaluated. Non-canonical splice sites were confirmed using reference proteins and transcripts. Rare erroneous reference annotations were corrected using TAIR10 annotations (https://www.arabidopsis.org/download/[34]. Genes were flagged with 'pseudogene' if a pseudogene from Araport11 was aligned to the same region. Incomplete genes and uncorrectable annotations were flagged. Genes at contig borders were flagged as 'truncated' if confirmed by protein or transcript mappings. Rarely, genes were extensively changed to rescue domain structures. These genes were flagged with 'mod'. Wrong gene models due to misassembled contigs were detected. Genes were flagged with 'misassembly' if base calls were contradicted reliably by CCS read mappings. Manual re-annotation was necessary to secure the reliability of NLR gene models.

#### Identification of paired NLRs

We generated a list of paired NLRs containing the nine Col-0 divergently transcribed TNLs sharing a genetic arrangement similar to the RPS4/RRS1 pair [35]. We added seven additional divergently transcribed pairs identified by manual inspection of 138 Col-0 genes that contained a TIR domain. We also used a CNL clone list (Dangl lab, unpublished) to mine the Col-0 genome for consecutive genes and included six paired CNL-CNL loci, of which only two are divergently transcribed. During manual curation we further identified one divergently transcribed pair of TNLs with no Col-0 allele and included it, too.

#### Identification of sensor-executor pairs

To further examine pair evolution, we made a narrower list of pairs. These are in head to head genetic orientation (in either the Col-0 reference genome, or in the assembled contigs where these gene pairs exist) and phylogenetically in the clades containing either RPS4 or SOC3 executor TNLs or the clades containing RRS1 or CHS1 sensor TN(L)s. The NB domain alignment-based phylogeny used to make this decision is presented in Supplementary Fig. 10. There are 16 such pairs identified in the pan-NLRome, two of which do not appear in the Col-0 reference genome. As a control group to test the possibility that genetic proximity could lead to co-evolution or conservation of population genetic characteristics, we identified a set of control pairs. These are pairs of NLR-encoding genes that are less than 4 kb apart in the Col-0 reference genome, but are not classified as sensor/executor pairs. We identified 15 such pairs, a list of all pairs specific to this context and pertaining to Figure 7b (main text) is provided in Supplementary Table 12.

#### Classification

Each gene that contained an NB, a TIR, an RPW8 domain, or a combination of those was defined as an NLR. The presence of LRR or CC motifs alone did not suffice our criteria. As a first subdivision (Supplementary Table 15), we defined TNLs (at least a TIR domain), CNLs (CC+NB domain), RNLs (at least a RPW8 domain), and NLs (at least a NB domain). The second subdivision defined 26 different groups by the different combinations of TIR, CC, NB, RPW8, LRR, and X (other Integrated Domains (ID)) independent of arrangement and number.

As mentioned earlier (see section: Web Apollo), protein domains were predicted using Pfam hmms. Coiled-coil (CC) motifs were refined in NLR genes using a majority vote from different prediction programs. In order to secure the correct annotation, Coils (2.2.1; InterProScan-defaults [36]), Paircoil2 (defaults [37]), and the NLR-parser (v.2; defaults [38]) predictions were compared to each other. Coils and Paircoil2 use databases of many known coiled-coils, whereas the NLR-parser uses two NLR-specific coiled-coil motifs (motif16 and motif17) [38]. CC signatures were considered credible if overlapping predictions existed in at least two of the three methods. Notably, CCs of functional NLRs previously published as CNLs are not always confirmed (Supplementary Table 16).

#### Architectures

An architecture is defined as the collapsed protein domain set in an NLR-coding gene. Domains occurring multiple times are reported only once in the reported architectures. Domains and collapsed architectures are reported in Supplementary Table 2 and Supplementary Table 9. The pair ratio reported in Fig. 2 was calculated by dividing the number of manually flagged paired NLRs by the total number of NLRs sharing the same architecture. A high order domain composition classification distinguishes between canonical and non-canonical domain architectures. Canonical architectures are strictly composed by any combination of NB (Pfam accession PF00931), TIR (PF01582), RPW8 (PF05659), LRR (PF00560, PF07725, PF13306, PF13855), or the Coiled-Coil structural motifs (Supplementary Fig. 2). Non-canonical architectures contain at least one ID, as defined in [39]. To be able to compare the pan-NLRome to the reference Col-0 accession we did not include the Col-0 RenSeq dataset in the architecture analyses. In order to identify novel and recurring domain arrangements, we compared the reference Araport11 Col-0 NLRs, with the pan-NLRome and the NLRome of 19 *Brassicaceae* species (Supplementary Table 6). In all domain architecture comparisons, we explicitly excluded the Col-0 Renseq gene models to enable the comparison to the reference Col-0, and included the *Brassicaceae* sets, whenever required.

Bash and R scripts for used to generate Fig. 2 UpSet plots, barplots and dotplots are available in github (see Fig. 2 script at https://github.com/weigelworld/pan-nlrome).

#### pan-NLRome Generation

The pan-NLRome of *A.thaliana* was constructed using a protein-clustering approach. Each protein cluster contained a set of homology-related NLRs from different ecotypes, and was termed an 'orthogroup' (Supplementary Table 17). Singletons were proteins that did not cluster with any other protein. Protein clusters were generated with a three step procedure. All-against-all full length protein alignments were produced (DIAMOND; version 0.9.1.102; --max-target-seqs 13169 --more-sensitive --comp-based-stats [40]). Putative ortholog and inparalog relationships were identified (orthAgogue, commit 82dcb7aeb67c, --use_scores --strict_coorthologs [41]). Protein clusters were formed based on the orthology information (mcl; version 12-135; -I 1.5 [42]).

##### Orthogroup Refinement

The initial set of orthogroups was inspected for over-clustering by screening for paralogs within orthogroups (Supplementary Fig. 11). A protein alignment for each orthogroup with more than 4 members was generated (T-Coffee; version 11.00.8cbe486; mode: mcoffee [43]). Protein sequence alignments were converted into the corresponding codon alignments (PAL2NAL; version 14, defaults [44]). The resulting codon alignment was used to remove three different types of outliers, namely non-homologous, partly mistranslated and low similarity sequences (OD-seq; version 1.0; --analysis bootstrap [45]). The remaining core sequences for each orthogroup were realigned in protein space. The protein alignments were converted into the corresponding codon alignments and used to infer a phylogenetic tree (FastME; version 2.1.5.1; -s -n -b 100 [46]). Each tree was used to detect simple paralogs (duplications in terminal branches) and complex paralogs (duplications spread across the whole phylogeny). For orthogroups where less than, or equal to 5% of its ecotype members showed duplications, all paralogs were removed. Otherwise the tree was split at (ecotype) duplication events (ete3; version 3.0.0b36 [47]) and new orthogroups were created from the leaves of all resulting sub trees (Supplementary Table 22).

Codon alignments and trees were re-computed as stated above (see alignments and trees at https://github.com/weigelworld/pan-nlrome).

A few misannotated genes were detected in OGs and removed from the NLRome (see Supplementary Table 24).

#### Orthogroup Classification

The final set of refined orthogroups was annotated with metadata derived from transcript-based majority votes (e.g., classes), transcript-based counts (e.g., members with IDs, members flagged as paired, members flagged as clustered) or orthogroup-based counts and analysis (e.g., type, diversity statistics, positive selection, average tree branch length). Refined orthogroups were classified into three size-based categories after visual inspection of the orthogroup size density distribution (Fig. 3a). Orthogroups with less than 13 members were typed as "cloud", orthogroups with more than 51 members "core", and those in between were typed "shell". We further classified orthogroups using protein domain architectures. We assigned a class and subclass to each orthogroup by using the majority vote from its members domain architectures.

Diversity and neutrality statistics were calculated for each codon alignment of the refined orthogroups (PopGenome; version 2.2.4 [48]). Domain-specific diversity statistics were calculated on subseted, concatenated alignments only consisting of positions covering the respective domains (e.g., NB). Alignment columns were annotated with a majority vote across all individual sequence annotations and selected subsequently. The average tree-derived branch length for an orthogroup was defined as the sum of all branch lengths normalized by the orthogroup size. Positive selection tests were carried out using HyPhy (version 2.3.13 [49]) using codon alignments and corresponding trees. Pervasive diversifying positive selection was detected with FUBAR (version 2.1; default parameters [50]) and sites considered with a posterior probability >= 0.95 (Supplementary Table 18 and Supplementary Table 19). Episodic diversifying positive selection was detected with MEME (version 2.0.1; default parameters [51]) and sites considered with a p-value threshold <= 0.01 (Supplementary Table 20 and Supplementary Table 21). Completely invariable codons were identified using a custom script (see *msa2cns* script at https://github.com/weigelworld/pan-nlrome). Domain-specific positive selection was calculated on a subset of positions covering the respective domains (e.g., NB). The alignment annotation was the same as for the Domain-specific diversity statistics.

An average expression percentage was estimated for each orthogroup using RNA-Seq data from the 1001 Genomes collection [3,52]. For each accession, a pseudo-transcriptome was generated from the accession-specific NLR transcripts and all non-NLR transcripts from the reference Col-0. NLR genes introns were added to the pseudo-transcriptome for expression filtering. Transcript abundance was quantified with pseudoalignments of RNA-Seq reads from 727 accessions of the 1001 Genomes collection (kallisto, v.0.43.0, --single -l 200 -s 25 -b 100 --bias [53]). The data was further processed with R (v.3.4.1). Abundance was normalized (DESeq2; v.1.16.1; estimateSizeFactor [54]) and expressed NLRs were defined using a per-accession expression threshold. Expression counts from introns were used to compute a background expression density distribution and subtracted from the density distribution of all NLR expression counts. The lowest expression level with a density > 0 was used as minimum expression threshold. On average, NLRs were considered expressed with an expected count >= 175. Finally, for each NLR, the percentage of accessions that provided reliable expression was calculated. Furthermore, we consulted the AtGenExpress (http://jsp.weigelworld.org/expviz/expviz.jsp) expression atlas to gauge absolute expression level [55], bias in leaf vs root specificity of expression and the pathogen inducibility of Col-0 NLRs. NLR genes were broadly divided into low, medium and high expression groups, based on whether at least two samples had absolute signal values in the developmental data sets that were 20 < expression < 100, 100 < expression < 1,000, 1,000 < expression. Genes that had generally absolute signal values below 20 were characterized as marginally expressed. If average expression in leaf and rosette samples was at least twice of that in root samples, or vice versa, genes were considered tissue biased in expression. Note that differences between tissues can be much larger, exceeding 100 fold. Pathogen inducibility was assessed from the AtGenExpress pathogen data set, based on consistent induction by at least two pathogen-related stimuli. The final Col-0 NLR annotation was amended to the respective orthogroups.

#### Orthogroup co-occurrence / Non-reference OG placement

Annotated non-NLR proteins in the 65 accessions were clustered into orthogroups using the same protein-clustering approach applied to NLRs above (see the ‘pan-NLRome Generation’ section). Briefly, non-NLR protein clusters were generated after three main procedures. First, we used DIAMOND to obtain all-against-all full length protein alignments (DIAMOND; version 0.9.1.102; --max-target-seqs 50691 --more-sensitive --comp-based-stats [40]. Second, we identified putative orthologs using orthAgogue (orthAgogue, commit 82dcb7aeb67c, --use_scores --strict_coorthologs [41]. Third, we used the MCL algorithm to define the cluster structure of the similarity relationships established in the previous steps (mcl; version 12-135; -I 1.5 [42]. No refinement steps were applied to non-NLR orthogroups (Supplementary Table 25).

For each NLR in NLR-OGs, we tested contig linkage with other annotated genes in the respective accession. To establish OG-OG co-occurrence, we extracted OG size (node size), NLR- and non-NLR-OGs (node color). Whenever OGs contained a Col-0 allele we established a reference anchoring position in the reference genome (node shape). OG co-occurrence matrices were used to calculate bidirectional networks showing contig linkage (edges). In the manuscript we report a minimum threshold of ten OG-OG co-occurrences for any anchoring/placement. OG Co-occurrence subnetworks containing successfully anchored/placed non-reference OGs were identified in Cytoscape v.3.5.1 [56], running on Java v. 1.8.0_151. To quantify OG-OG co-occurrences in the selected subnetworks, we extracted all observed combinations from the accessions gff files and visualized co-occurrence intersections in UpSet plots. Putative paired NLRs were identified by testing OG enrichment in annotation flags (see 'Paired NLRs' and 'Over-represented flags analyses' methods section). Flag enrichment was calculated using Fisher and hypergeometric tests and FDR (Supplementary Table 23). All enrichments with a q-value below 0.1 were reported. Finally, the anchoring positions of anchored OGs is shown as a schematic karyogram (see 'Figure Generation' methods section). Reference Col-0 NLR coordinates were extracted from the TAIR9/Araport11 annotation. Equivalence between Col-0 Araport11 and RenSeq 6909 NLR identifiers was by reciprocal best blast hit. Non-reference OG anchoring positions are approximate values derived from manual inspection of NLRome assemblies (see 'Web Apollo' methods section). Bash-, R-scripts and input files used to generate the UpSet plots, karyogram and the cytoscape network are available in github (see Fig. 4 script at https://github.com/weigelworld/pan-nlrome).

#### Saturation Analysis

Orthogroup as well as haplotype and nucleotide diversity discovery rates were determined by saturation analysis. For orthogroup discovery, accessions were randomly selected from the pan-NLRome and the number of orthogroups counted they were part of. The process was repeated

1,000 times starting with two and ending with 64 randomly selected accessions. For nucleotide and haplotype diversity discovery, accessions were selected as above mentioned. Nucleotide and haplotype diversity were calculated for each of the replicates and average. The process was repeated 100 times starting with two and ending with 64 randomly selected accessions.

##### Orthogroup Visualization

Raw orthogroups and corresponding metadata were integrated in iTOL [57] for visualization and reinspection (https://itol.embl.de/shared/pan_NLRome). Protein trees showed the evolutionary history of each orthogroup. Similarity between members was also reflected in branch lengths and bootstrap values. The multiple sequence alignment showed sequence variation on base pair level. The identifiers of refined orthogroups were added to show over-clustered orthogroups and outliers. The domain architecture and the protein length were plotted to compare orthogroup members structurally. Transposable elements (TEs) are known to influence gene activity or be the cause of gene duplication. TEs in exons, introns, and 2kb up- or downstream of NLRs were integrated in iTOL. Sub-clustering might be related to accession-based metadata, thus we included for each protein if its accession belonged to the relict group, the geographic origin, and the admixture group.

##### Over-represented flags analyses

Gene annotation flags in each orthogroup were compiled using an in house bash script. Flag enrichment was calculated in R using hypergeometric test. Multiple testing was corrected via false discovery rate (FDR) estimation and q-values below 0.1 were reported (Supplementary Table 23).

#### Figure Generation

Bash and R scripts for used to generate Fig. 2 UpSet plots, barplots and dotplots are available in github (see Fig. 2 script at https://github.com/weigelworld/pan-nlrome). Bash, R scripts, input files and cytoscape network are available in github (see Fig. 4 script at https://github.com/weigelworld/pan-nlrome).

All quantitative figure panels were generated using R (version 3.4.4 [58]) and RStudio [59], unless otherwise stated. For clarity, floating text was added to SVG files generated in R using Inkscape (version 0.92.3 https://inkscape.org). Used packages included ggplot2, grid, gridExtra, reshape2, gsubfn, cowplot, rworldmap, yarrr, UpSetR, PerformanceAnalytics, karyoploteR and viridis. OG phylogenetic trees were visualized using iTOL [57]. The phylogeny for Supplementary Fig. 10. was generated through the use of MEGA 6.06 including MUSCLE [60] and WAG maximum-likelihood phylogenetics [61]. Fig. 7c and Supplementary Fig. 10 were visualized using FigTree (v1.4.3; https://github.com/rambaut/figtree). Input data and R scripts for all relevant figures can be found at https://github.com/weigelworld/pan-nlrome.

Supplementary Figure 1.

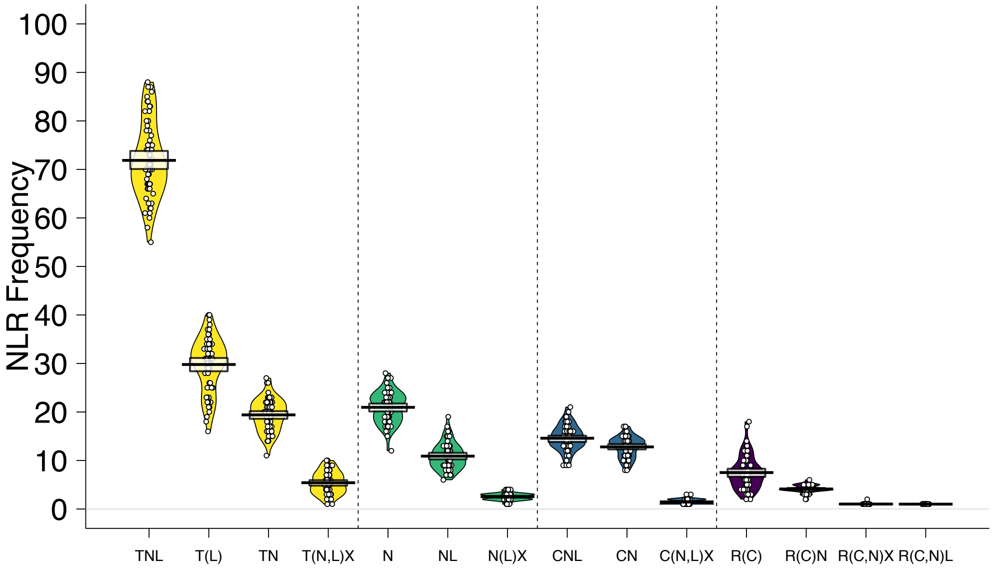


**Supplementary Figure 1. NLR frequency for different subclasses.** For each subclass, the corresponding class is color coded (TNLs: yellow, NLs: green, CNLs: blue, and RNLs: purple), and classes are in addition divided by the vertical dashed lines. NLRs are grouped into subclasses by their domains content: T (TIR), N (NB), C (CC), R (RPW8), and X (all other integrated domains). Each domain must be present at least once, domains in brackets may be present. Domain order is not considered. The mean is shown as a solid black horizontal line and the 95% Highest density Intervals (HDI: points in the interval have a higher probability than points outside) are shown as solid bands around the sample mean. All raw data points are plotted as open circles and the full densities are shown as a bean plot.

### Supplementary Figure 2.

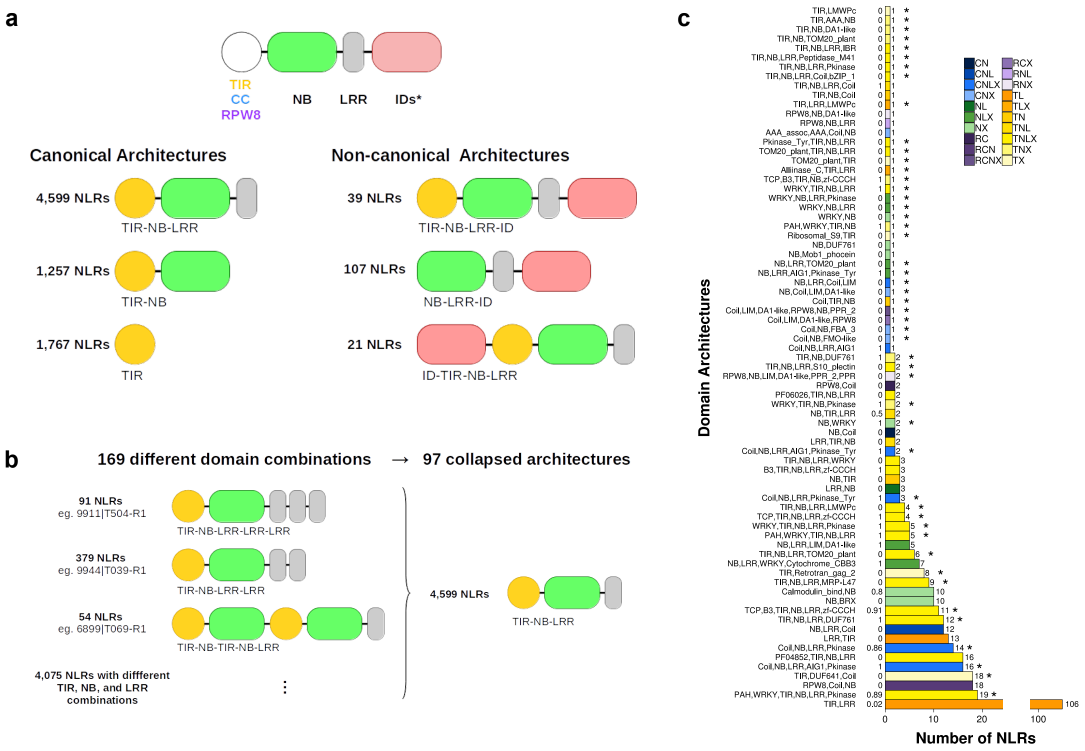


**Supplementary Figure 2. Schematic representation of NLR domain architecture diversity and simplification of consecutively repeated domains.** a) Examples of NLR domain architecture diversity. On top, a generic NLR, with an ID (Integrated Domain) is shown at the C-terminus. IDs can also be found at the N-terminus, and more rarely between the three canonical domain types. b) Reduction of domain combinations by collapsing duplicated/repetitive domains. The number of NLRs grouped by each of the original architectures is shown on the left, along with one example that can be visualized in the genome browser. Ellipsis in the bottom left represent 19 other architectures containing 4,079 proteins exclusively composed of TIR, NB and LRR domains. The same strategy was applied to all other architectures containing at least one duplicated domain in the RPW8, NB and CC classes. c) Full set of the novel A. thaliana NLR architectures. Includes the architectures contributed by only one gene. Domain architectures are shown in the y-axis. The number of NLRs in each architecture is shown in the x-axis. Asterisks indicate the 49 architectures not yet detected in the Brassicaceae outside of A. thaliana, or in the reference accession Col-0. Numbers next to y-axis show the ratio of paired NLRs divided by the total number of NLRs in each architecture.

### Supplementary Figure 3.

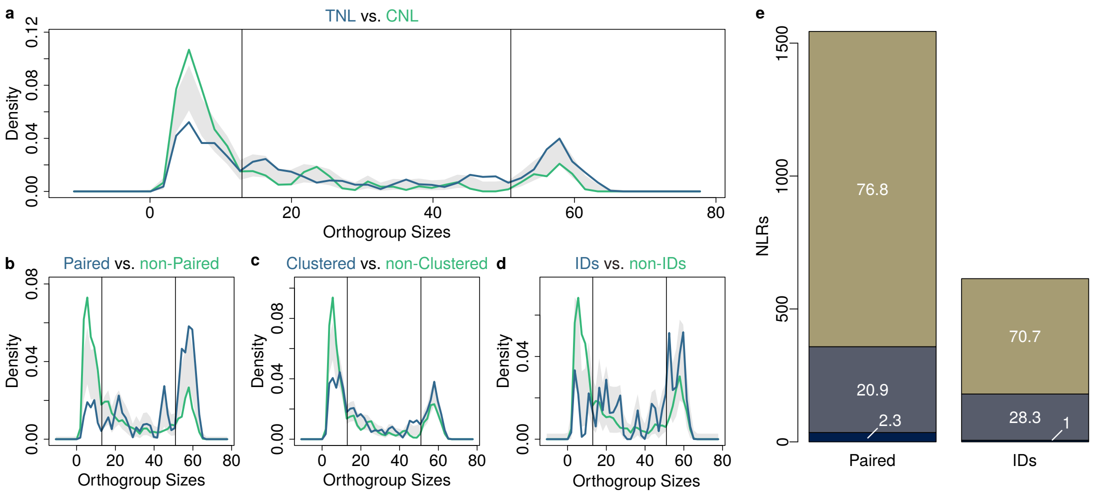


**Supplementary Figure 3. OG size distribution comparisons.** Vertical black lines divide cloud (left section) from shell (middle section) and core (right section) NLRs. a) Comparison of OG size distributions of TNL OGs (blue) and CNL OGs (green) b) Comparison of paired (blue) and non-paired (green) OGs. c) Comparison of clustered (blue) and non-clustered (green) OGs. d) Comparison of ID-containing (blue) and non-ID-containing OGs (green). e) Distribution of Paired NLRs and NLRs with IDs across the Cloud- (dark blue), the Shell- (grey), and the Core- (olive green) pan-NLRome.

### Supplementary Figure 4.

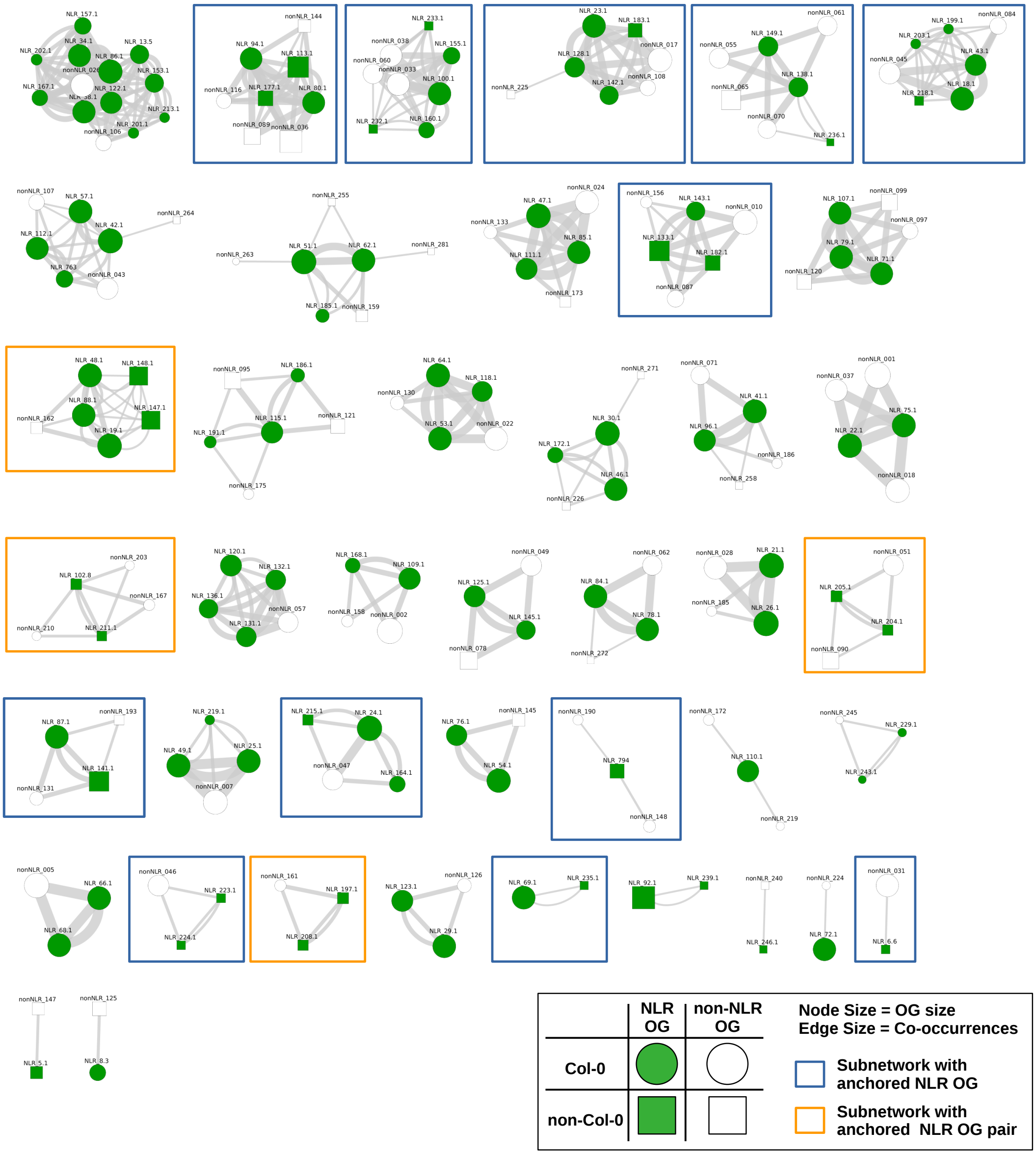


**Supplementary Figure 4. Orthogroup co-occurrence network.** Annotated NLR (green nodes) and non-NLR genes (white nodes) clustered into OGs were analyzed for co-occurrence in the same contig. The number of co-occurrences is represented by grey lines connecting nodes (edges). The minimal co-occurrence threshold imposed was 10 accessions, but similar networks can be derived for any number accessions. NLR OGs without a Col-0 allele (green square nodes) are highlighted in blue boxes. Hypothetically paired OGs not known in Col-0 are highlighted in orange boxes.

Supplementary Figure 5.

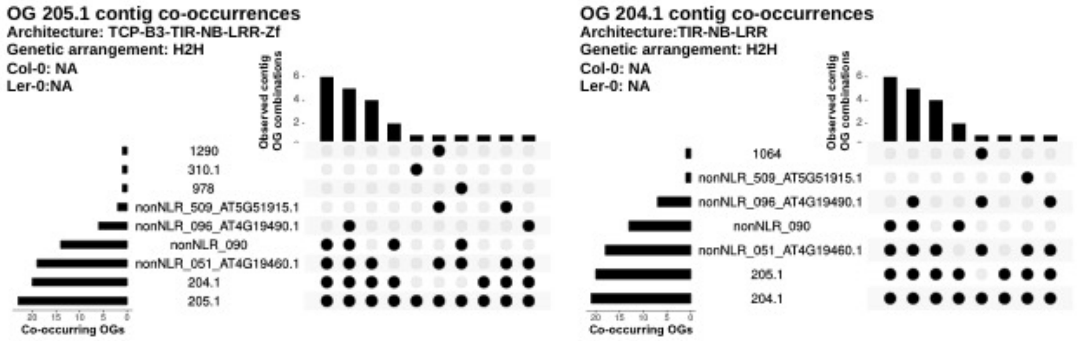


**Supplementary Figure 5. Quantitative co-occurrence of the novel hypothetical paired NLRs in OG205.1 and OG204.1.** Abbreviations: OG, Orthogroup; H2H, Head-to-head; NA, Not available.

### Supplementary Figure 6.

a

b

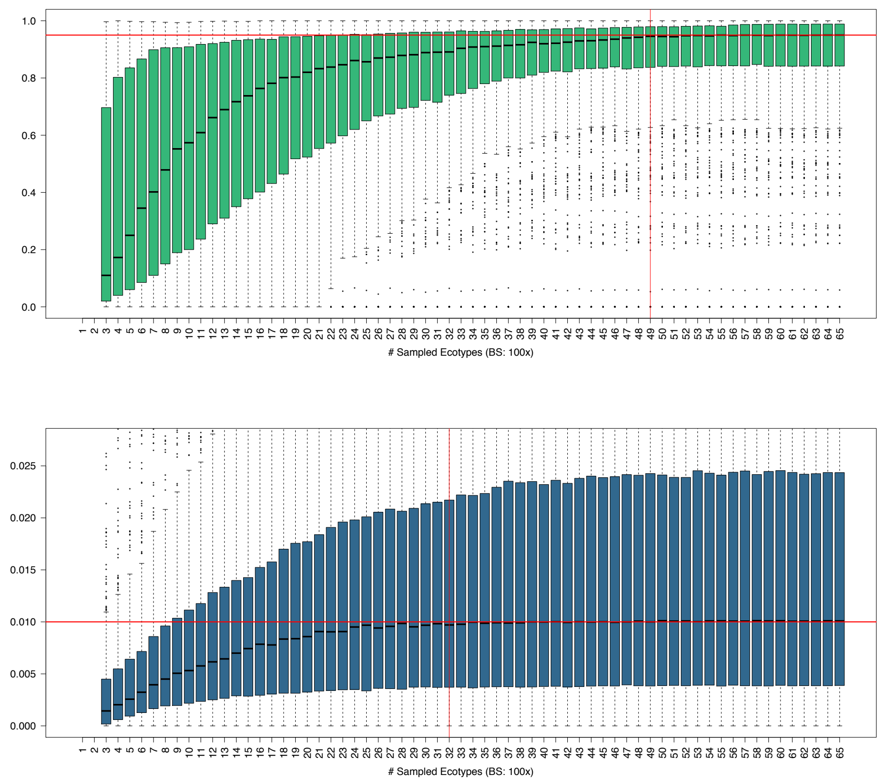


**Supplementary Figure 6. Nucleotide and haplotype diversity saturation.** Saturation of nucleotide (a) and haplotype (b) diversity after random subsetting of the complete Pan-NLRome into bins of increasing sizes. For each size 100 bootstrap were carried out.

Supplementary Figure 7.

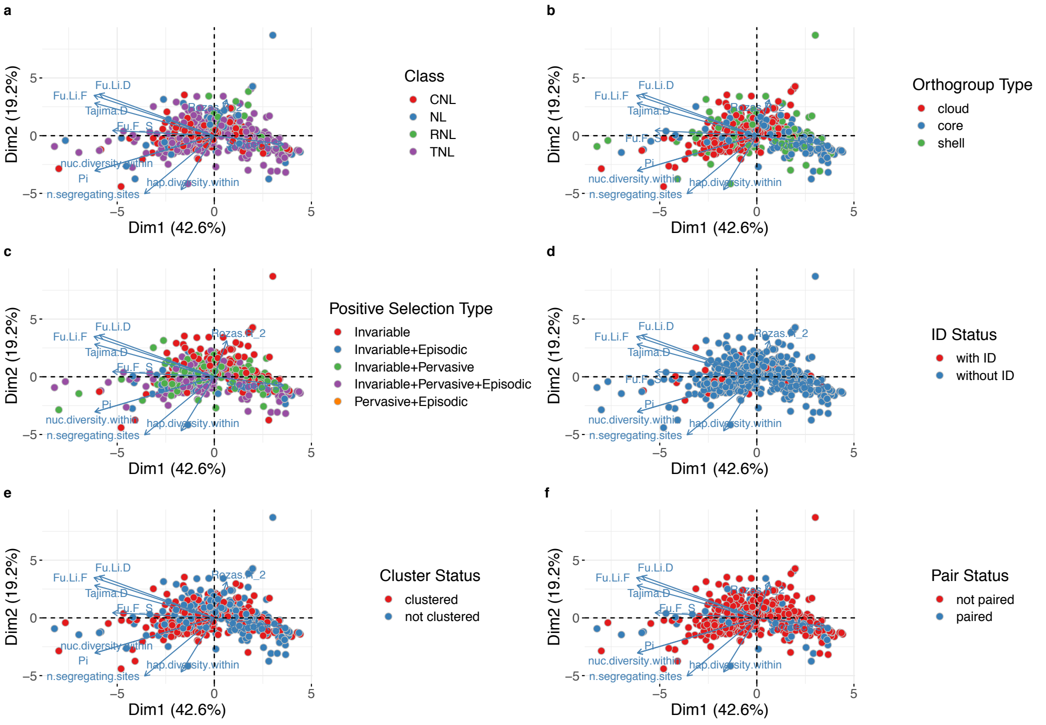


**Supplementary Figure 7. Population genetics statistics based PCA.** Principal component analysis carried out on 10 population genetics statistics, namely Nucleotide diversity / Pi, Haplotype diversity, Fu and Li's D, Fu and Li's F, Tajima's D, Rozas' R2, Strobeck's S and the number of segregating sites. Panels are colored according to categorical variables.

### Supplementary Figure 8.

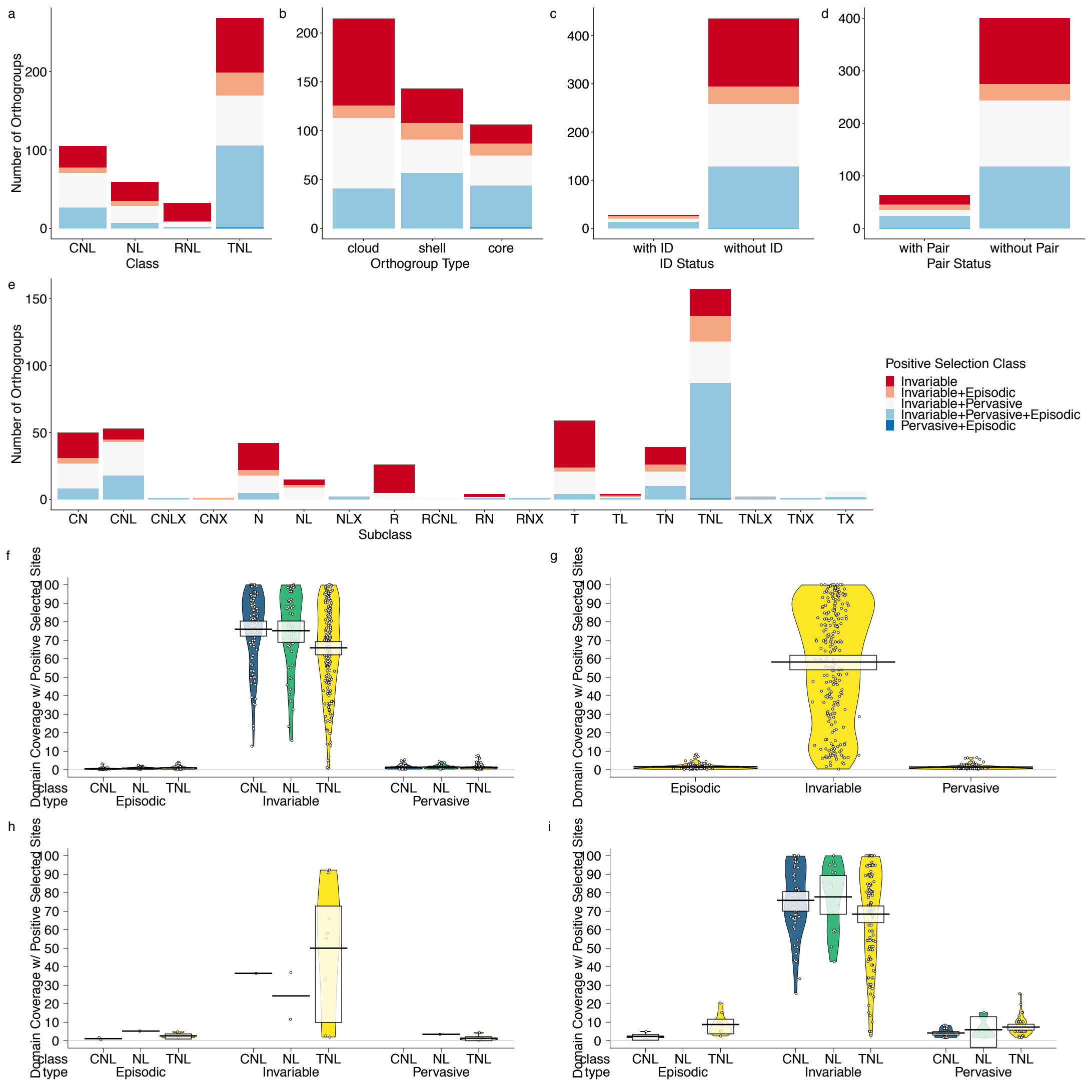


**Supplementary Figure 8. Positive selection landscape of the Pan-NLRome.** (a-e) Absolute number of orthogroups in positive selection classes grouped by NLR class (a), orthogroup type (b), presence of a non-canonical domain (c), presence of a paired NLR (d) or NLR subclasses (e). An orthogroup was considered if at least one positive selected site of a given class was detectable. (f-i) Domain coverage with positively selected sites grouped by NLR class and positive selection type across canonical domains (f-g, i) and all aggregated non-canonical domains (h).

### Supplementary Figure 9.

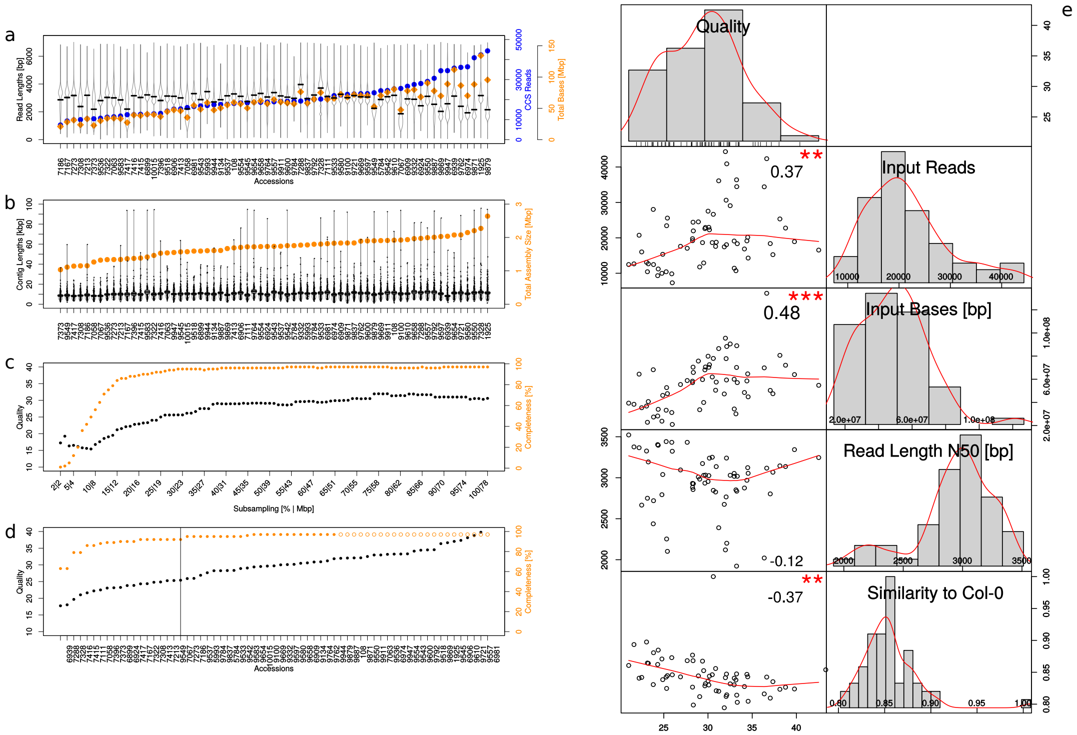


**Supplementary Figure 9. Read and assembly statistics.** a) Read lengths distribution (Q20-filtered CCS reads) for all accessions (black). The mean is shown as a solid black horizontal line. The full densities are shown as a bean plot. The total number of CCS reads (blue circles) and the total number of bases (orange diamonds) are plotted in addition. b) Contig lengths distribution (black). The mean is shown as a solid black horizontal line and the 95% Highest density Intervals (HDI: points in the interval have a higher probability than points outside) are shown as solid bands around the sample mean. The full densities are shown as a bean plot. Raw data points are plotted using black dots. The total assembly sizes (orange circles) are plotted in addition. c) Quality (black) and completeness values (orange) for sub-sampled Col-0 datasets. The amount of input data for each sub-sampling experiment is shown as a second x axis. d) Quality (black) and completeness values (orange) for all RenSeq accessions. Unfilled circles indicate accessions with qualities higher than any sub-sampled dataset. The vertical black line is drawn at 95% completeness. e) Correlations between the Assembly Quality, the amount of Input Reads, the amount of Input Bases [bp], the read length N50 [bp], and the similarity to Col-0 are shown for the RenSeq datasets. Histograms and kernel densities (red line) are plotted for each variable. Scatter plots for variable pairs are shown together with a fitted line (red) and the Pearson's correlation coefficient (significance 0 '***',0.001 '**',1 ' ').

### Supplementary Figure 10.

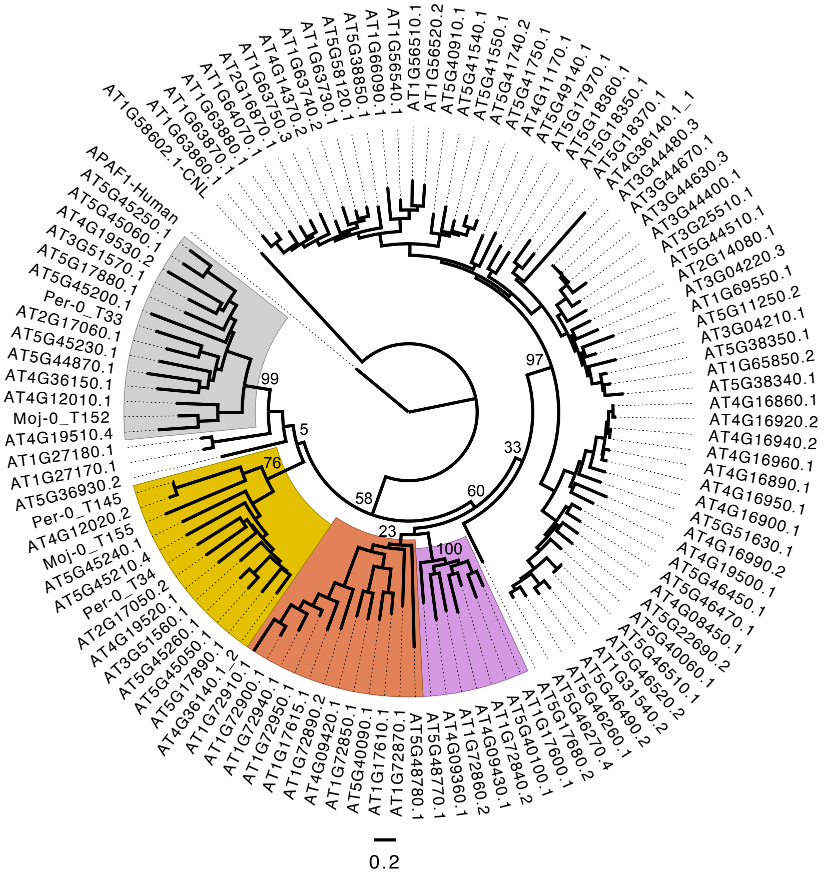


**Supplementary Figure 10. Phylogenetic tree of NB domain alignments for refining sensor/executor pairs from all pairs.** RPS4/RRS1-like and SOC3/CHS1-like paired TNLs fall into distinct subclades. These are indicated by color: RPS4-like (Silver, executors), RRS1-like (Gold, sensors), SOC3-like (Pink, executors) and CHS1-like (Bronze, sensors). This phylogeny was constructed by aligning the NB domain (~240 amino acids) of all TIR and NB containing Col-0 proteins and selected additional representatives of pair flagged orthogroups (OGs) from the pan-NLRome that are not represented in Col-0 (identified by their OG and protein numbers). NB domains from APAF1 (Human) and AT1G58602.1 (A. thaliana CNL) were also included. Amino acid sequences were aligned with MUSCLE (Neighbor joining clustering), refined by manual trimming and the phylogeny produced with the WAG maximum likelihood method allowing for 3 discrete Gamma categories. AT4G36140 contains two distinct NB domains, both of which were included and the second of which clusters with other RRS1-like NB domains. Number of 100 bootstraps supporting topology shown at major node vertices. Scale bar represents amino acid substitutions per site.

### Supplementary Figure 11.

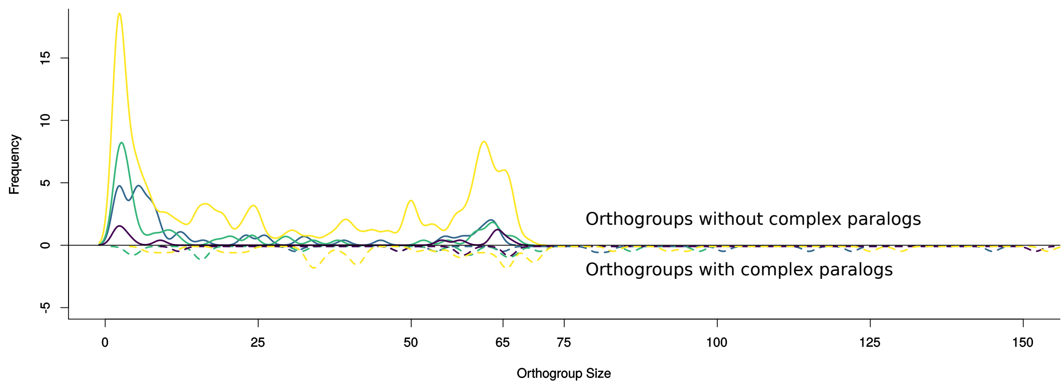


**Supplementary Figure 11. Orthogroup (OG) size frequencies before OG refinement.** Data are shown separately for the different NLR classes (TNL: yellow, NL: green, CNL: blue, RNL: purple) and for OGs with (solid lines) and without (dashed lines) complex paralogs (duplications spread across the whole phylogeny).
